# Glioblastoma–natural killer cell crosstalk: insights from dynamic spheroid models reveal the importance of secreted cytokines and the CD155 axis

**DOI:** 10.1101/2025.10.29.685263

**Authors:** Anamarija Habič, Tina Kolenc Milavec, Pia Žižek, Špela Kladnik, Bernarda Majc, Emanuela Senjor, Milica Perišić Nanut, Andrej Porčnik, Borut Prestor, Urban Švajger, Metka Novak, Barbara Breznik

**Author notes:** **Corresponding authors:** Barbara Breznik, Anamarija Habič.

## Abstract

Glioblastoma (GB) is an aggressive primary brain cancer with poor patient prognosis. Natural killer (NK) cells can recognise and eliminate a range of malignant cells, including GB stem cells, which drive GB recurrence. NK cell-based immunotherapy has emerged as a promising approach for GB treatment, but a better understanding of the complex crosstalk between GB and NK cells is needed, particularly within the immunosuppressive GB tumour microenvironment. In this study, we established a reproducible protocol for the production and dynamic culture of uniformly sized GB spheroids using the Celvivo Clinostar system. Our spheroids recapitulated the heterogeneous structure of GB and expressed ligands for NK cell receptors at levels distinct from those observed in corresponding GB cell lines in standard culture, implicating altered sensitivity of GB cells to NK cells in dynamic 3D cultures. GB-NK cell crosstalk was GB cell type dependent and the ability of NK cells to infiltrate GB did not necessarily correlate with their cytotoxicity against GB cells. Spheroids derived from differentiated GB cells secreted higher levels of immunomodulatory cytokines compared to spheroids from GB stem-like cells, and a prominent increase in the secretion of immune-attracting factors was observed in their co-cultures with NK cells. Finally, the CD155-DNAM1/TIGIT axis was indicated as an important regulator of NK cell cytotoxicity against GB stem-like cells. Collectively, our results highlight important factors in GB-NK cell communication and provide a groundwork for further targeted research as well as therapeutic evaluation of NK cell-based approaches in the established dynamic 3D cultures.

## 1. Background

Glioblastoma (GB) is the most common and aggressive primary brain cancer in adults. Despite the standard of care treatment that includes surgery, radiotherapy and chemotherapy with temozolomide, the prognosis of GB patients is poor and less than 10% of patients are still alive at 5 years after the diagnosis (1,2). During the past decades, there have been no major advances in GB treatment which would significantly improve patient survival, indicating an urgent need for novel therapeutic approaches.

In the last decade, immunotherapy has revolutionized the treatment of many cancers, especially haematological malignancies (3). Nevertheless, applying these approaches to solid tumours has often shown limited success. To a great extent, this can be attributed to the immunosuppressive tumour microenvironment (TME), which not only represents a physical barrier for immune cell infiltration, but also directly regulates the activity of immune cells by secreting immunomodulating and often immunosuppressive factors (4). Despite several challenges such as intracranial location of the tumour, immunosuppressive TME, and heterogeneity of GB, immunotherapy holds promise for GB treatment (5).

Several immunotherapeutic approaches have already been tested in the setting of GB (5). However, although a number of pre-clinical studies showed encouraging results, to date, no durable clinical benefits have been observed in randomized clinical trials in patients. This indicates a lack of relevant preclinical models that would reliably reflect the complexity and biological properties of the GB in human and could accurately predict the efficacy of immunotherapies (6). Unlike 2D cell cultures, 3D culture models, such as spheroids, organotypic cultures, and organoids, more closely mimic tumour architecture, nutrient and oxygen gradients and metabolism and thus allow for a more accurate assessment of immunotherapeutic approaches (7,8). Nevertheless, the absence of a complete TME, systemic immunity and functional vasculature still limit the predictive power of these models. (Humanized) immunocompetent mouse models of GB, although more relevant in these regards, are expensive and time-consuming and often do not fully reproduce human pathology and immune responses (9). To an extent, blood flow and immune cell infiltration may be mimicked in vitro by culturing human cancer cells in static or perfused microfluidic platforms (10).

Natural killer (NK)-cell based immunotherapeutic approaches may be particularly feasible in the setting of GB. NK cells are innate lymphoid cells that can eliminate tumour cells without prior sensitization and recognition of tumour-associated antigens (11). Their activity is regulated by activating and inhibitory receptors, which interact with ligands on target cells. If the interactions with activating receptors prevail over the inhibitory signals, the intracellular cell cytotoxic machinery is activated, leading to elimination of the target cells (11,12). Most commonly, NK cells operate through the perforin-granzyme pathway, releasing cytolytic granules and inducing target cell apoptosis. Alternatively, target cells can also be eliminated through the induction of the death receptor pathway (13,14). Another important mechanism employed by NK cells which links the innate and adaptive immune responses and can also be exploited therapeutically is the antibody dependent cellular cytotoxicity (ADCC) (15). The term relates to the ability of NK cells to eliminate antibody-coated target cells, which are recognised by the Fc region-binding receptor CD16 (FcγRIII) on the surface of NK cells. Lastly, NK cells also secrete a variety of cytokines such as interferon-gamma (IFN-γ) and tumour necrosis factor-alpha (TNF-α), which directly affect cancer cells and modulate both innate and adaptive immune responses (11,12).

As GB is characterised by a low tumour mutational burden (16), the use of NK cells may be beneficial compared to T cell-based therapies and vaccine-based approaches to bypass the lack of tumour-associated antigens and also the lack of functional antigen-presenting cells in the highly immunosuppressive GB TME. Additionally, NK cells may represent a powerful tool against the pool of heterogeneous GB cells, as (unlike T cells) they can recognize a broad spectrum of target cells (17,18). Importantly, NK cells have been shown to kill GB stem cells – a subset of cancer cells that are intrinsically resistant to a number of standard therapeutic approaches and represent key drivers of GB recurrence (19,20). As no graft-versus-host disease is induced by NK cells, allogeneic off-the-shelf NK cell products may be used for patient treatment, providing a cheaper, more scalable and reproducible alternative to patient-specific T cell products. Due to their immediate availability, off-the-shelf products are particularly convenient for the treatment of aggressive types of cancer (17).

In the GB TME, the abundance of NK cells is generally low, accounting for approximately 2–10% of leukocytes within the tumour (21), but the presence of activated NK cells positively correlates with patient survival (22). However, in the immunosuppressive TME, fuelled by GB cells, the infiltrated NK cells often exhibit altered phenotypes and restrained cytotoxic activity. Several immunosuppressive factors in the GB TME were reported to downregulate the expression of activating NK receptors and decrease NK cell secretion of IFN-γ (23–25). Additionally, NK cell number and function may also be impaired by hypoxia (26).

A deeper understanding of the GB-NK cell crosstalk is needed to develop effective NK cell-based therapeutic approaches for GB and prevent or revert its potential resistance mechanisms. Relevant in vitro models should be used to investigate the GB-NK cell interplay and NK cell-based therapeutic approaches in the human setting. To address these needs, we established a reproducible approach for generation and dynamic culture of GB spheroids from GB cell lines in the Celvivo Clinostar system. GB spheroids were co-cultured with NK cells in direct co-culture and on a dynamic organ-on-chip platform to model and explore the GB-NK cell crosstalk. Moreover, the importance of the CD155 axis for the cytotoxicity of NK cells was studied in more detail.

## 2. Methods

### Cell lines and culture

NIB140 is a differentiated GB cell line derived from patient tumour tissue. Tumour tissue was obtained from University Medical Centre Ljubljana. The study was approved by the National Medical Ethics Committee of the Republic of Slovenia (approval No. 0120-190/2018-2711-41). NIB140 cells were cultured in surface-treated tissue culture flasks (Jet Biofil) in high-glucose Dulbecco’s modified Eagle’s medium (DMEM; Gibco, Thermo Fisher Scientific) supplemented with 10% foetal bovine serum (FBS; Gibco, Thermo Fisher Scientific) and 1× penicillin/streptomycin (Sigma-Aldrich). Prior to reaching confluency, cells were regularly passaged by 0.125% trypsin–EDTA solution (Gibco, Thermo Fisher Scientific).

GB stem-like cell line NCH421k was purchased from Cell Lines Service GmbH (CLS). Cells were cultured as floating spheres in suspension cell culture flasks (Sarstedt) in Neurobasal Medium (NBE; Gibco, Thermo Fisher Scientific) supplemented with 1× penicillin/streptomycin (Sigma-Aldrich), 2 mM L-glutamine (Sigma-Aldrich), 1× B-27 (Invitrogen), 1 U/mL heparin (Sigma-Aldrich), 20 ng/mL basic fibroblast growth factor (Invitrogen) and 20 ng/mL epidermal growth factor (Invitrogen). Spheres were regularly dissociated with TrypLE Express (Gibco, Thermo Fisher Scientific).

Cell line NK-92 was purchased from the American Type Culture Collection (ATCC). Cells were grown in NK-92 complete medium, i.e., RPMI 1640 medium with GlutaMAX (Gibco, Thermo Fisher Scientific) supplemented with 12.5% FBS (Gibco, Thermo Fisher Scientific), 12.5% heat inactivated horse serum (Gibco, Thermo Fisher Scientific), 1× penicillin/streptomycin (Sigma-Aldrich) and 200 IU/mL recombinant human interleukin 2 (IL-2; Miltenyi Biotec). For activation, cells were grown for 24 h in NK-92 complete medium supplemented with 1,000 IU/mL recombinant human IL-2 (Miltenyi Biotec).

Isolated healthy donor peripheral blood mononuclear cells (PBMCs) were obtained as buffy coats from the Blood Transfusion Centre of Slovenia. Experiments were performed in accordance with the approval of National Medical Ethics Committee of the Republic of Slovenia (approval no. 0120-279/2017-3). Cells were washed twice in PBS with 1 mM EDTA and 2% FBS to remove platelets. Healthy donor NK cells were isolated from PBMCs using the EasySep Human NK Cell Enrichment Kit (Stem Cell Technologies) and the EasySep cell separation magnet (Stem Cell Technologies) according to the manufacturer’s instructions. For isolation of NK cells from whole blood of GB patients, PBMCs were isolated using the Lympholyte® Cell Separation Media (Cedarlane) and NK cells were isolated as described above. Experiments were performed in accordance with the approval of National Medical Ethics Committee of the Republic of Slovenia (approval no. 0120-190/2018-2711-41). After isolation, NK cells were grown in NK complete medium, i.e., RPMI 1640 medium with GlutaMAX (Gibco, Thermo Fisher Scientific) supplemented with 12.5% FBS (Gibco, Thermo Fisher Scientific) and 1× penicillin/streptomycin (Sigma-Aldrich). For activation, cells were grown for 18h in NK complete medium supplemented with 1,000 IU/mL recombinant human IL-2 (Miltenyi Biotec).

Normal human astrocytes (NHA) were purchased from Lonza and were cultured in Astrocyte Medium (Sciencell) supplemented with 10% FBS (Gibco, Thermo Fisher Scientific), 1% astrocyte growth supplement (ScienCell), and 1× penicillin/streptomycin (Sigma-Aldrich). Prior to reaching confluency, cells were passaged by 0.125% trypsin–EDTA solution (Gibco, Thermo Fisher Scientific). Low passages were used in experiments.

K562 cells were purchased from the American Type Culture Collection (ATCC) and were grown in RPMI 1640 medium with GlutaMAX (Gibco, Thermo Fisher Scientific) supplemented with 10% FBS (Gibco, Thermo Fisher Scientific) and 1× penicillin/streptomycin (Sigma-Aldrich). Cells were passaged every 2-3 days.

Cell lines were regularly checked for Mycoplasma contamination using the MycoAlert Mycoplasma Detection Kit (Lonza).

### Immunophenotyping of PBMCs and NK cells isolated from PBMCs

8-Color Immunophenotyping Kit, anti-human (Miltenyi Biotec), was used according to the manufacturer’s instructions to determine the frequencies of immune cell populations (T cells, B cells, NK cells, monocytes, neutrophils, eosinophils, CD4+, CD8+, and CD56+CD3+ T cell subsets) in PBMCs. The kit was also used to assess the purity of the NK cells isolated from PBMCs. Stained cells were analysed using the MACSQuant Analyzer 10 Flow Cytometer (Miltenyi Biotech) and FlowJo software (BD Life Sciences). The purity of NK cells isolated from PBMCs was > 90%.

### Establishment and culture of GB spheroids in the Celvivo Clinostar system

GB spheroids were established by the forced floating method in 96-well round bottom microplates (Falcon, Corning). The starting numbers of cells per spheroid were optimized for each cell line and were set at 5,000 and 30,000 cells for NCH421k and NIB140, respectively. The selected number of cells was seeded in 100 ul of corresponding complete medium with 4% methylcellulose. Plates were centrifuged for 90 minutes at 900 g and incubated for 3 days under static conditions at 37 °C in 5% CO_2_ atmosphere. Spheroids were then resuspended in fresh complete media, transferred to pre-equilibrated ClinoReactors (Celvivo) and cultured in a dynamic system in the ClinoStar clinostat incubator (Celvivo) at 37 °C and 5% CO_2_ atmosphere. The speed of ClinoReactor rotation was adjusted daily to ensure optimal spheroid dispersion. Media in the ClinoReactors were changed every 2–3 days. Spheroids that had been grown in ClinoReactors for 8 days were used for experiments.

### Monitoring spheroid growth

Spheroid growth was monitored by measuring the spheroid area. Images of spheroids in ClinoReactors were taken on Nikon Eclipse Ts2R inverted microscope. Spheroid area was quantified in Fiji (27).

### Dissociation of GB spheroids into single-cell solution

For flow cytometry and calcein release assay spheroids were dissociated into single cells. NCH421k spheroids were dissociated in TrypLE Express (Gibco, Thermo Fisher Scientific). After 5 minutes of incubation at room temperature, spheroids were dissociated by pipetting. Cells were washed with PBS and resuspended in desired staining solution/medium. For dissociation of NIB140 spheroids, we used TrypLE Express (Gibco, Thermo Fisher Scientific) and Colagenase, Type I (Thermo Fisher Scientific) diluted in DMEM (Gibco, Thermo Fisher Scientific). Spheroids were incubated in the enzyme mixture at 37 °C and were occasionally mixed by pipetting until they dissociated. Single cell suspension was washed with PBS and resuspended in desired staining solution/medium.

### Flow-cytometric determination of cell viability and death in GB spheroids

After spheroid dissociation, cells were incubated with annexinV-FITC (Miltenyi Biotec) for 15 min at 4°C. Then, cells were diluted and stained for 5 min with 7-AAD (Miltenyi Biotec). Stained cells were diluted and analysed using the MACSQuant Analyzer 10 Flow Cytometer (Miltenyi Biotech) and FlowJo software (BD Life Sciences).

### Preparation and immunofluorescent staining of formalin-fixed, paraffin-embedded (FFPE) sections

For preparation of the FFPE sections, spheroids (without or with infiltrated NK cells) were washed with PBS and fixed in buffered 4% formaldehyde solution (Merck) for 72 h at 4 °C. Fixed samples were embedded in 0.5% agarose in PBS to ease further processing and prevent sample loss. Agarose blocks were dehydrated and embedded in paraffin. FFPE sections (4 μm thick) were prepared for subsequent immunofluorescent staining.

FFPE sections were deparaffinised in xylene (Chem-Lab) and rehydrated in a series of ethanol solutions of decreasing concentrations (100%, 96%, and 70%). Samples were heated at 95 °C for 20 min in 10 mM sodium citrate buffer (pH 6.0) to retrieve antigens. Blocking was performed for 1 h at room temperature in 10% (v/v) normal goat serum (Sigma-Aldrich), 0.1% Triton X-100 (v/v) (Sigma-Aldrich) and 1% bovine serum albumin (BSA; w/v) (Sigma-Aldrich) in PBS. Samples were incubated with primary antibodies diluted in 1% BSA (w/v) in PBS (Table S1) overnight at 4 °C. After washing with 0.5% BSA (w/v) in PBS, fluorescently labelled secondary antibodies diluted in 1% BSA (w/v) in PBS (Table S1) were added to the slides and incubated for 1 h at room temperature. The slides were then washed in PBS and nuclei were stained with Hoechst 33258 solution (Sigma-Aldrich) diluted 1:1,000 in PBS. After washing with PBS, samples were mounted in ProLong Gold AntiFade reagent (Invitrogen, Life Technologies), cover slipped and sealed with nail polish. Inverted fluorescent microscope (Nikon Eclipse Ti, Tokyo, Japan) and NIS-Elements, Nikon software, were used to image fluorescence.

### Flow cytometry to assess the expression of receptors on NK cells and their ligands on cancer cells

The expression of receptors on live NK cells (CD16, CD96, DNAM-1, KIR2DL1, KIR2DL2/L3/S2, KIR2DL4, KIR3DL1, NKG2D, NKp30, PD-1, TIGIT) was measured on NK-92 cells, NK cells isolated from PBMCs of 6-9 healthy donors and NK cells isolated from PBMCs of 6 GB patients. The same receptors were also analysed on NK-92 cells that have been cultured for 48 h in a 1:1 mixture of NK-92 complete medium and NCH421k/NIB140 conditioned medium. The expression of NK cell receptor ligands (B7-H6, CD112, CD155, PD-L1, CD54, HLA-A,B,C, HLA-E, MICA/MICB, ULBP-1, ULBP-2/5/6, ULBP-3) was measured on live flask- and spheroid-cultured cancer cell lines NCH421k and NIB140.

Cells were resuspended in PBS with 1% BSA and stained with antibodies listed in Table S2. IgG isotypic controls were used as controls. Cells were stained for 30 min at 4°C. After washing, cells labelled with fluorescently conjugated primary antibodies were analysed by the MACSQuant Analyzer 10 Flow Cytometer (Miltenyi Biotech). In the case of unconjugated primary antibodies, cells were washed and additionally stained with Alexa Fluor 488 rabbit anti-mouse IgG secondary antibody (Thermo Fisher Scientific), incubated for 30 min at 4°C, washed and analysed on the MACSQuant Analyzer 10 Flow Cytometer (Miltenyi Biotech). FlowJo software (BD Life Sciences) was used to determine the % of cells positive for specific receptor or ligand. Gates were set based on the corresponding IgG isotypic control samples.

### Establishment of direct NK cell co-cultures with GB spheroids

Spheroids that had been grown in ClinoReactors for 8 days were used for establishment of co-cultures with NK cells (NK-92 or healthy donor NK cells). At least 5 spheroids of each cell line were dissociated to determine average number of cells per spheroid and to calculate the number of (non-)activated NK cells needed for the selected effector:target ratios (2.5:1 or 5:1). For the co-cultures, spheroids with 100 µl of corresponding complete media were transferred to wells of a round bottom 96-well plate (Falcon, Corning). The concentrations of (non-)activated NK cells were adjusted to achieve the desired final effector:target ratios after the addition of 100 µl of NK cell suspension per spheroid in 96-well plate. For activated NK cells, IL-2 was included in the media to achieve final concentration of 1,000 IU/mL. Spheroids and NK cells were co-cultured for 24 h at 37 °C in 5% CO_2_ atmosphere.

For the analysis of NK cell infiltration and cytotoxicity by flow cytometry, NK cells were fluorescently labelled prior to the onset of co-cultures to enable their direct detection and discrimination. NK cells were washed with PBS and stained with 5 µM CellTracker Blue CMAC (Thermo Fisher Scientific) or 10 µM CellTracker Green CMFDA (Thermo Fisher Scientific) in RPMI medium without supplements. After 30 minutes of incubation, cells were washed, and their concentration was adjusted to achieve the desired final effector:target ratios after the addition of 100 µl of NK cell suspension per spheroid in 96-well plate.

Co-cultures of GB spheroids and nonactivated or IL-2-activated PBMCs from GB patients were established in the same manner.

### Flow cytometric analysis of NK cell spheroid infiltration and cytotoxicity

Spheroids with infiltrated fluorescently pre-labelled NK cells were washed in PBS and individually dissociated as described above. Cells were washed and resuspended in 100 µl PBS with 1% BSA. 1 µl 7-AAD (Miltenyi Biotec) was added and incubated for 5 min to label dead cells. Cells were diluted and analysed by the MACSQuant Analyzer 10 Flow Cytometer (Miltenyi Biotech) and FlowJo software (BD Life Sciences) to determine NK cell infiltration and their cytotoxicity against GB cells.

### Analysis of cytokines, chemokines, growth factors and cytotoxicity-related factors in the media of co-cultures

After the 24 h incubation, media of the co-cultures were collected for the analysis of secreted cytokines, chemokines, growth factors and cytotoxicity-related factors. Media were centrifuged (5 min, 5,000 rpm) and supernatants were stored at −80 °C. Cytokine/chemokine analysis was performed in two multiplex assays (Human Cytokine/Chemokine 96-Plex Discovery Assay® Array (HD96) and TGFB 3-Plex Discovery Assay® Multi Species Array (TGFβ1-3)) by the Eve Technologies Corporation (Calgary, AB Canada) on the Luminex® 200™ platform. The following targets were analysed: BAFF, CCL1, CCL13, CCL17, CCL19, CCL2, CCL20, CCL21, CCL22, CCL23, CCL24, CCL26, CCL27, CCL28, CCL3, CCL4, CCL5, CCL7, CCL8, CXCL10, CXCL11, CXCL13, CXCL16, CXCL5, CXCL9, EGF, Eotaxin, FGF-2, FLT-3L, Fractalkine, GCP-2, G-CSF, GM-CSF, Granzyme A, Granzyme B, GROα, HMGB1, IFN-α2, IFNβ, IFNγ, IFN⍰, IL-10, IL-11, IL-12p40, IL-12p70, IL-13, IL-15, IL-16, IL-17A, IL-17E/IL-25, IL-17F, IL-18, IL-1RA, IL-1α, IL-1β, IL-2, IL-20, IL-21, IL-22, IL-23, IL-24, IL-27, IL-28A, IL-29, IL-3, IL-31, IL-33, IL-34, IL-35, IL-4, IL-5, IL-6, IL-7, IL-8, IL-9, LIF, Lymphotactin, M-CSF, MIP-1⍰, PDGF-AA, PDGF-AB/BB, Perforin, sCD137, sCD40L, SCF, SDF-1, sFas, sFasL, TGF-α, TGF-β1, TGF-β2, TGF-β3, TNFSF13, TNFα, TNFβ, TPO, TRAIL, TSLP, and VEGF-A.

### Analysing NK-92 cell infiltration in a dynamic organ-on-chip platform

We set up a dynamic MIVO® Single-Organ Platform (MIVO® platform, React4life) – a commercially available setup, consisting of transwell chambers carrying spheroids in Matrigel (Corning) under flow conditions, mimicking the influx of immune cells into the tumours. NCH421k or NIB140 spheroids were prepared by the forced floating method (as described above) and embedded in Matrigel (Corning) in the tumour chamber above a microcirculation of NK-92 cells. NK-92 cells at effector:target ratio 2.5:1 were applied to the circulation at flow rate 1 mL/min for 24 h at 37 °C in 5% CO_2_ atmosphere. IL-2 was included in the media to achieve final concentration of 200 IU/mL. NK-92 cells were pre-labelled with 10 µM CellTracker Green CMFDA (Thermo Fisher Scientific) or 5 µM CellTracker Blue CMAC (Thermo Fisher Scientific) in RPMI medium without supplements. After 24 h of NK-92 cell circulation, contents of the tumour chamber were washed with cold PBS to remove Matrigel. Spheroids were dissociated and cells were resuspended in 2% FBS in PBS. Dead cells were stained by 7-AAD (Miltenyi Biotec). MACSQuant Analyzer 10 Flow Cytometer (Miltenyi Biotech) and FlowJo software (BD Life Sciences) were used to determine the infiltration of labelled NK cells into the tumour chambers and spheroids and the viability/death of GB and NK-92 cells.

### Calcein release assay

Calcein release assay was used to test cytotoxicity of (IL-2-preactivated) NK-92 and healthy donor (HD) NK cells against NCH421k and NIB140 cells from standard or spheroid cultures. It was also used to test whether NK-92 and HD NK cell cytotoxicity against K562 cells is affected by pre-culturing NK cells in NCH421k or NIB140 conditioned medium.

Serial dilutions of effector cells (NK-92 or HD NK cells) in at least three technical replicates were prepared in a round bottom 96-well plate (Falcon, Corning) in final volume of 100 µl corresponding complete medium per well. Target cells were labelled with Calcein-AM (Sigma) at final concentration 15 µM in serum-free RPMI 1640 medium (Gibco, Thermo Fisher Scientific) for 30 min at 37 °C. After the incubation, cells were washed and resuspended in corresponding complete medium to a final concentration 0.1×10^6^ cells/mL. 5,000 labelled target cells (50 µl) per well were added to the effector cells and control wells. The plate was centrifuged at 200 g for 2 min and incubated at 37 °C and 5% CO_2_ for 3 h. After incubation, the plate was centrifuged at 700 g for 7 min and 50 µL of the supernatant from each well was transferred to a black 96-well plate (Thermo Fisher Scientific). Fluorescence (excitation: 496 nm, emission: 516 nm) was measured on a Synergy Mx microplate reader (Bio-Tek Instruments Inc.). Based on the calcein fluorescence measurements, the percentage of cytotoxicity was calculated as 100 × (test release – spontaneous release)/(total release – spontaneous release). Spontaneous release was measured in wells containing 100 µl NK-92 complete or NK complete medium and 50 µl calcein-AM-labelled target cells. For total release, 2% Triton X-100 was added to the NK-92 complete or NK complete medium to achieve complete lysis of calcein-AM-labelled target cells. Lytic units (LU) were calculated using the inverse of the number of effector cells needed to lyse 30% of the target cells multiplied by 100.

Calcein release assay with blocking of the CD155 axis was executed as described above with the addition of CD155 blocking antibodies (clone L95, Merck), DNAM-1 blocking antibodies (clone 102511, R&D Systems), TIGIT blocking antibodies (clone MBSA43, Thermo Fisher Scientific) or their combinations. Target cells (NCH421k or NIB140 cells) were preincubated for 45 min with CD155 blocking antibodies and NK cells were preincubated for 45 min with DNAM-1 or/and TIGIT blocking antibodies. All blocking antibodies were used at concentration of 10 ug/mL. Isotype control at the same concentrations was used as control.

### Statistical analysis

Unless stated otherwise, graphs were plotted and statistics calculated in GraphPad Prism version 10.4.1. At least three biological replicates (N≥3) were performed per experiment. Data are presented as mean ± standard error of the mean (SEM); sample sizes and statistics used are included in figure legends. p values⍰<⍰0.05 were considered to indicate significant differences. The p-values are indicated in the figures as follows: ^∗^p < 0.05; ^∗∗^p < 0.01; ^∗∗∗^p < 0.001 and ^∗∗∗∗^p < 0.0001.

Heatmap visualisation of secreted factors and hierarchical clustering based on the “one minus Pearson correlation” metric were performed in Morpheus, https://software.broadinstitute.org/morpheus.

## 3. Results

### 3.1 GB spheroids cultured in the dynamic Celvivo Clinostar system recapitulate the heterogeneous architecture of GB

We established a novel spheroid model derived from human GB cell lines which closely mimics the architecture of GB tissue (Fig. 1a). To our knowledge, we are the first to maintain GB spheroids in the dynamic Celvivo Clinostar system, which provides a controlled and physiologically relevant culture environment. By ensuring low shear stress and even distribution of nutrients, it enables long-term spheroid culture. In our study, spheroids were successfully prepared and maintained from three distinct GB cell lines: NCH421k, a suspension GB cell line with stem-like characteristics, NIB140, our in-house patient-derived adherent GB cell line, and U87, a commonly used adherent GB cell line (Fig. 1b, Fig. S1), indicating a general applicability of the approach. The production and culture of spheroids in ClinoReactors was reproducible and gave round and uniformly sized spheroids which could be cultured in the Celvivo Clinostar system for at least several weeks (Fig. S1).

**Figure 1.**
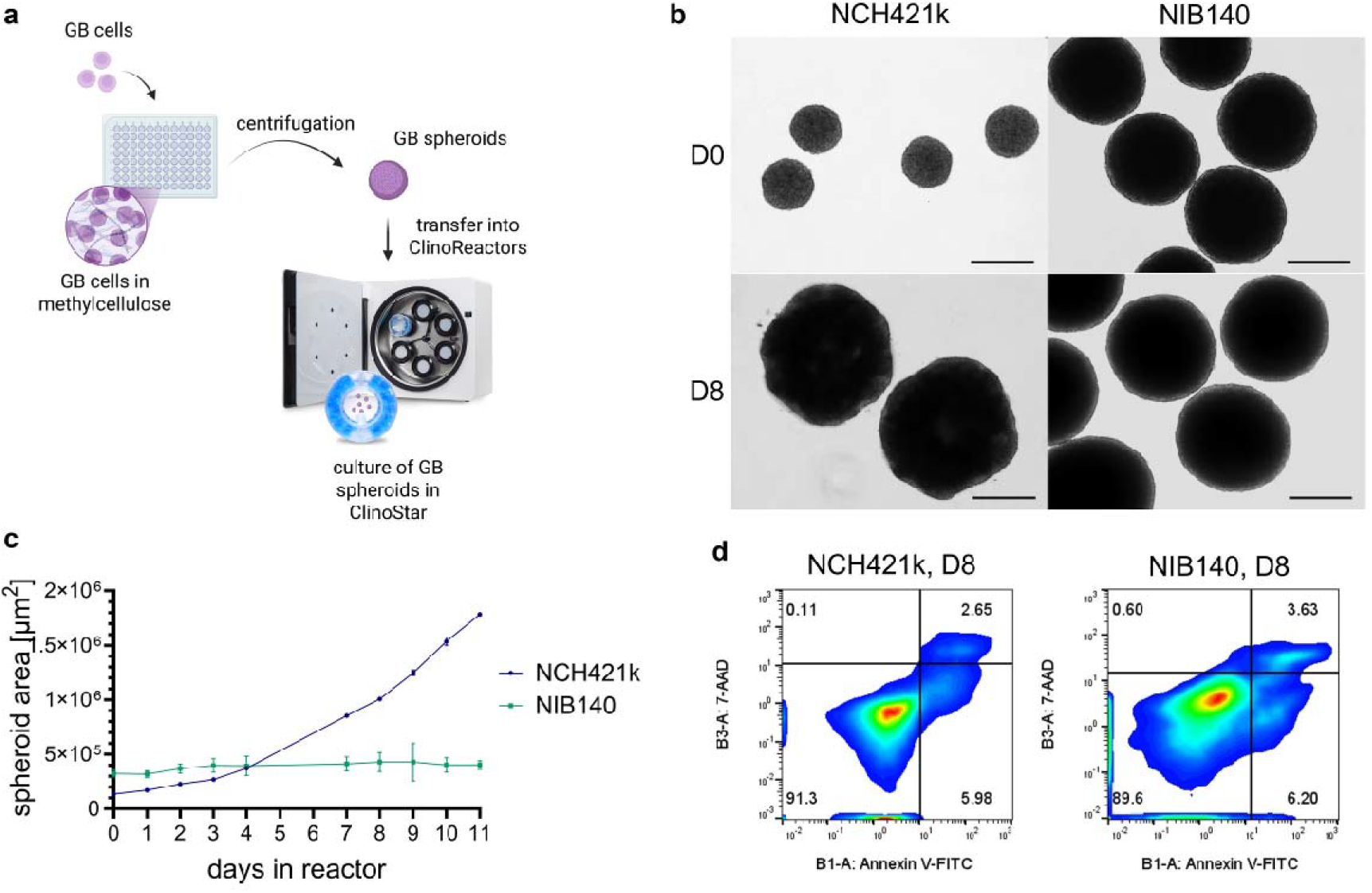
Growth of GB spheroids in the Celvivo Clinostar system. **a)** Schematic representation of the preparation and culture of GB spheroids established in this study. Created with bioRender.com. **b)** NCH421k and NIB140 spheroids in ClinoReactors on days 0 and 8. Scalebar: 500 µm. **c)** Growth curves of NCH421k and NIB140 spheroids. Growth was determined by measuring spheroid area in time. Mean ± SEM of at least three biological replicates is presented. **d)** Analysis of cell viability in NCH421k and NIB140 spheroids on day 8. In both spheroid types, the proportion of cells in early or late stages of apoptosis is below 10%.

Spheroids derived from cell lines NCH421k and NIB140 were characterized in more detail and were used for further experiments. Cell lines NCH421k and NIB140 exhibit major differences in expression of stemness-related genes, including *OLIG2, SOX2, PROM1, CD9*, and *NOTCH* (Fig. S2) and were used as proxies of GB stem-like and differentiated GB cells, respectively. Major differences were observed in growth kinetics of NCH421k and NIB140 spheroids. NCH421k spheroids grew much faster than the NIB140 spheroids (Fig. 1c, Fig. S1) and possessed a loose structure, while the NIB140 spheroids were more compact. On day 8 in the ClinoReactors, i.e., the timepoint at which spheroids were used for further experiments, approximately 10% of cells were detected in early or late stages of apoptosis in spheroids of both cell lines (Fig. 1d).

We further characterized GB spheroids at the molecular and spatial level by immunofluorescent staining of stemness markers SRY-box transcription factor 2 (SOX2) and oligodendrocyte transcription factor 2 (OLIG2), differentiation marker glial fibrillary acidic protein (GFAP), mesenchymal subtype marker CD44, proliferation marker protein Ki-67, and hypoxia markers hypoxia inducible factor 1 subunit alpha (HIF1α) and carbonic anhydrase 9 (CA9). The staining was performed at two different time points, i.e., after 8 days and 18 days of culture in the ClinoReactors (Fig. 2, Fig. S3). We confirmed the expected expression of GB cell markers and differences between the NCH421k and NIB140 cell lines (Fig. 2a). Low levels of differentiation marker GFAP and mesenchymal subtype marker CD44 were detected in NCH421k spheroids, while both were profoundly expressed in NIB140 spheroids. In NCH421k spheroids, practically all cells were positive for stem cell markers SOX2 and OLIG2, while no OLIG2 and lower levels of SOX2 were observed in NIB140 spheroids. Interestingly, the expression of SOX2 was not uniform among the NIB140 cells, indicating an intrinsic heterogeneity of cell states. In line with the slower growth rate of NIB140 spheroids compared to NCH421k spheroids, the abundance of Ki-67-positive cells was lower in the NIB140 spheroids. In spheroids that had been cultured in the ClinoReactors for 18 days, heterogeneous zones were observed, including regions of highly proliferative cells as well as some hypoxic and necrotic regions deeper in the spheroid structure (Fig. 2b). Thus, our GB spheroids recapitulated the spatially heterogeneous architecture of GB *in vivo*.

**Figure 2.**
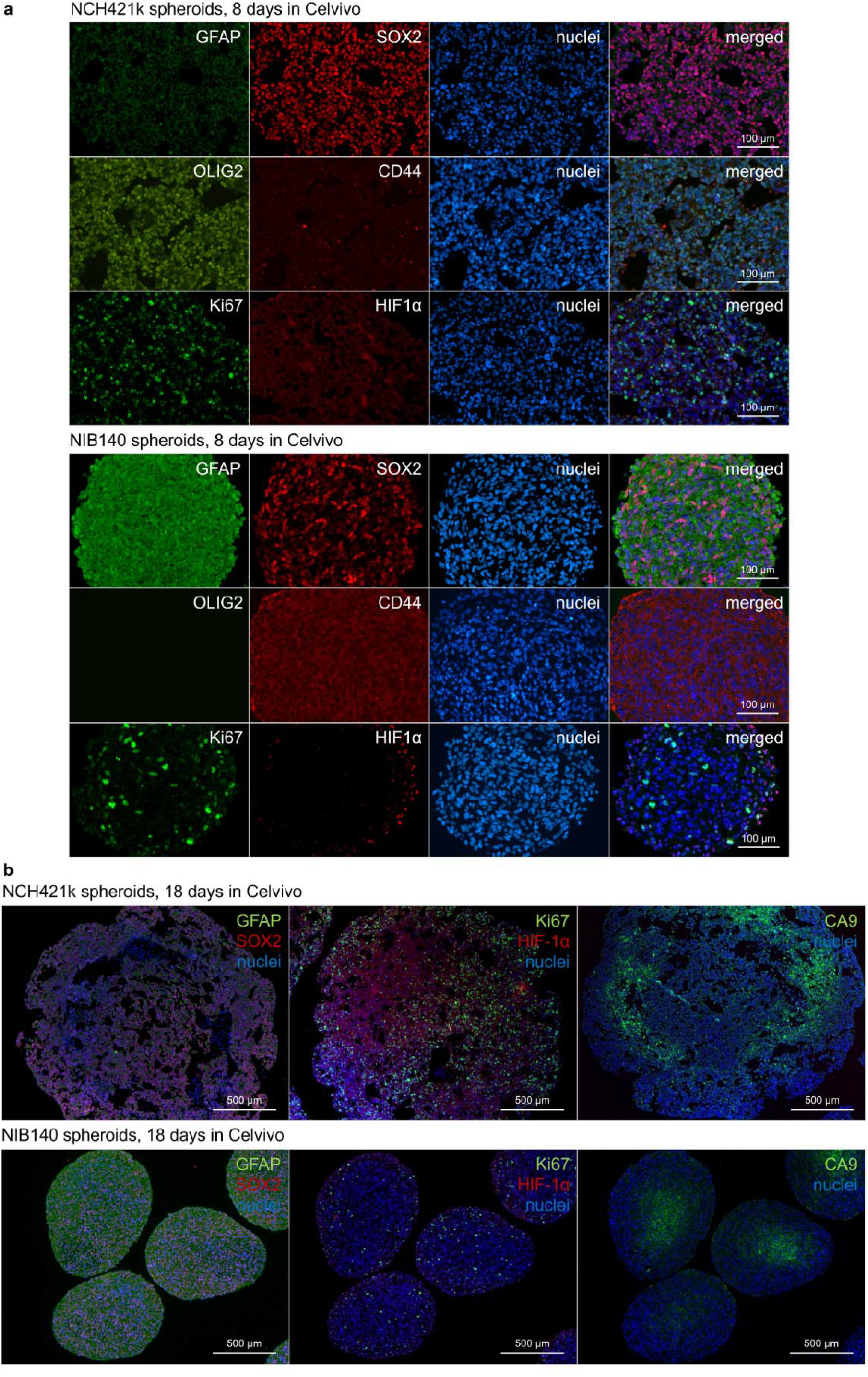
Characterisation of GB spheroids cultured in the Celvivo Clinostar system. **a)** Immunofluorescent staining of selected markers in NCH421k and NIB140 spheroids on day 8. NCH421k spheroids express high levels of SOX and OLIG2 (stem cell markers) and low levels of GFAP (differentiation marker) and CD44 (mesenchymal subtype marker). Spheroids are highly proliferative, as indicated by a high proportion of Ki-67-positive cells. Cells in the NIB140 spheroids express high levels of GFAP and CD44, no OLIG2 and varying levels of SOX2. The proportion of Ki-67-positive proliferative cells is lower compared to NCH421k spheroids. **b)** Immunofluorescent staining of selected markers in NCH421k and NIB140 spheroids on day 18. Heterogeneous zones were observed in both types of spheroids, such as regions of low and high cell proliferation, necrotic and hypoxic regions.

### 3.2 Spheroid culture affects the expression of ligands for NK cell receptors on GB cells and their sensitivity to NK cells

As we aimed to use our spheroid model to study the crosstalk between GB and NK cells, we analysed the expression of several ligands for NK cell receptors in GB cell spheroids and, in parallel, in GB cell lines from standard culture (Fig. 3a). The expression of poliovirus receptor (CD155), nectin-2 (CD112), B7 homolog 6 (B7-H6), intercellular adhesion molecule 1 (CD54/ICAM-1), UL16 binding proteins (ULBPs), MHC class I polypeptide-related sequences A and B (MICA-B), HLA class I histocompatibility antigen, alpha chain A/B/C and E (HLA-A/B/C and HLA-E), and programmed cell death receptor ligand 1 (PD-L1) was analysed. A number of NK cell-related ligands were prominently expressed on NCH421k and NIB140 cells and several differences were observed between the two cell lines in standard culture: CD155, B7-H6, CD54, ULBP1, and ULBP-2,5,6 were expressed at higher levels in NIB140 cells compared to NCH421k cells, which expressed more CD112. In spheroid cultures, NCH421k cells expressed more HLA-A/B/C and CD155 and less ULBP-2,5,6 and B7-H6 compared to the NIB140 cells.

**Figure 3.**
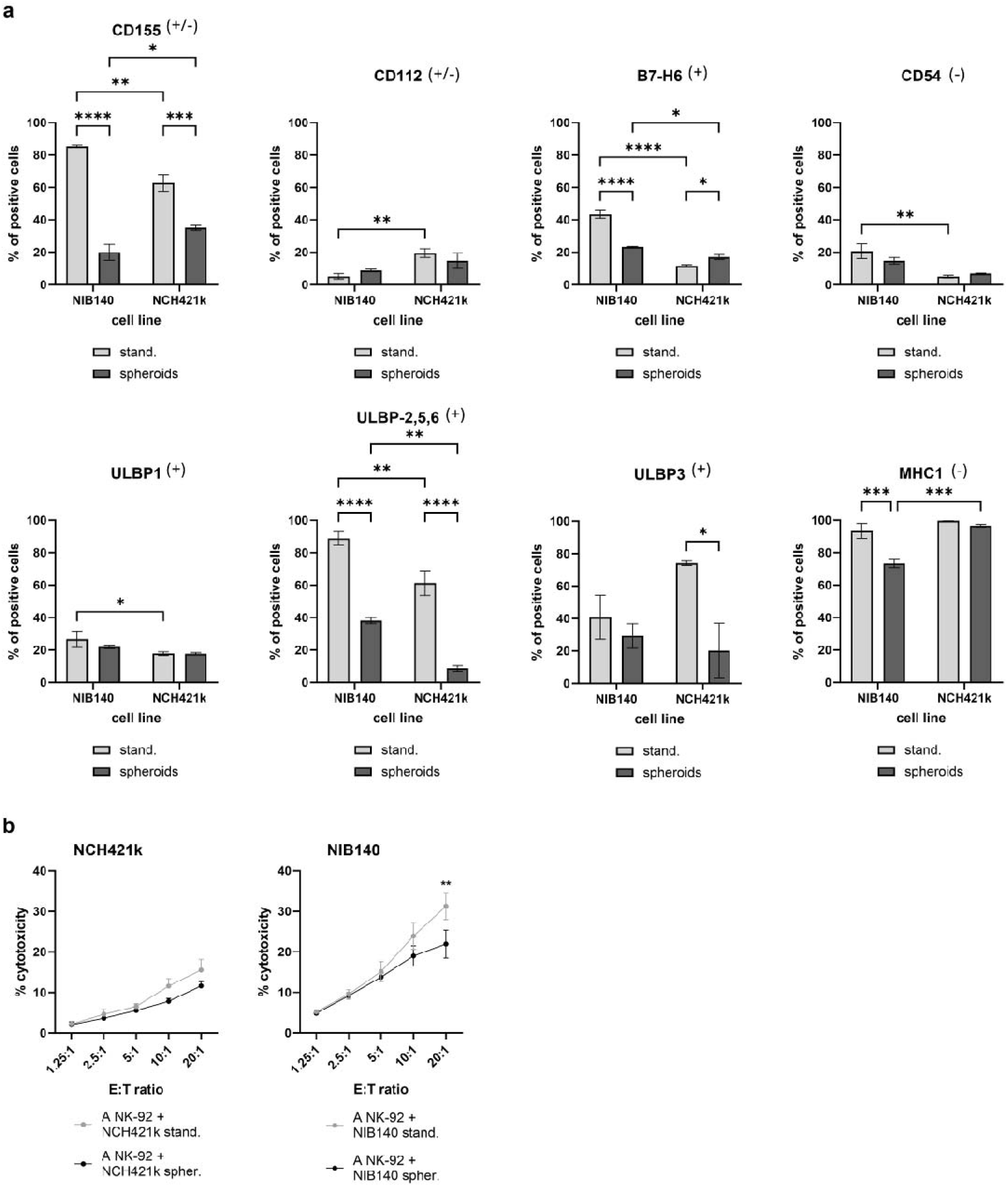
Spheroid culture affects the expression of ligands for NK cell receptors on GB cells and their sensitivity to NK cells. **a)** The expression of selected NK cell ligands was analysed by flow cytometry. For each of the two cell lines, the expression was also analysed on cells in standard culture. Data are presented as mean⍰±⍰SEM of three biological replicates. Two-way ANOVA with uncorrected Fisher’s least significant difference was used to calculate statistical significance of the differences in expression levels of the ligands between cell lines and culture methods. The symbols in parentheses designate ligands that bind activating (+), inhibitory (−) or either activating or inhibitory NK receptors (+/−). **b)** % cytotoxicity plots of calcein release assay of NK-92 cells against dissociated NCH421k and NIB140 spheroids and NCH421k and NIB140 cells from the standard culture. Data are presented as mean ±⍰SEM of 3–4 biological replicates. 2-way ANOVA with Šídák’s multiple comparisons test was used for statistical analysis.

The expression levels of several ligands were significantly affected by the culture method (standard vs. Celvivo spheroid culture) (Fig. 3a). For example, high percentages of CD155 positive cells were observed in standardly cultured NIB140 and NCH421k cultures (85% and 63%, respectively), while the proportion of CD155 positive cells was much lower in spheroid-cultured NIB140 and NCH421k cells (20% and 35%, respectively). A similar decrease in the percentage of ligand-positive cells in spheroids was observed for ULBP-2,5,6 (in both cell lines), B7-H6 (in NIB140 spheroids), HLA-A/B/C (in NIB140 spheroids) and ULBP3 (in NCH421k spheroids).

To further test whether GB cell sensitivity against NK cells is affected by the culture type, calcein-release assay was performed on single cells dissociated from GB Celvivo spheroids as well as on cells from standard culture of both cell lines (Fig. 3b, Fig. S4a-c). NK-92 cells exhibited slightly higher cytotoxicity against NCH421k and NIB140 cells from standard culture compared to cells from spheroids; NIB140 cells in spheroids showed significantly lower sensitivity to NK-92 cell cytotoxicity at the highest effector:target ratio (20:1) (Fig. 3b). Conversely, the calcein-release assay read-out showed no significant differences between cytotoxicity of HD NKs against cells from standard and spheroid culture (Fig. S4b-c). Activation of HD NKs with IL-2 increased their cytotoxicity, particularly in case of the NCH421k cells. In comparison, normal human astrocytes were resistant against NK-92 cells (Fig. S4d).

### 3.3 NK-92 cells show differences in infiltration rates between GB cell lines in direct co-cultures and in a dynamic organ-on-chip platform

To study how NK cells interact with GB cells in a 3D microenvironment, we set up GB spheroid-NK cell co-cultures. NK-92 and HD NK cells were applied to the GB spheroids from the Celvivo Clinostar system for 24 h. Then, co-cultures were dissociated to single cells or fixed for the preparation of FFPE slides and NK cell infiltration was measured by flow cytometry or immunofluorescence staining, respectively (Fig. 4a). NK-92 cells poorly infiltrated the NCH421k spheroids, while the infiltration was profound in the NIB140 spheroids (Fig. 4b). Nevertheless, infiltration levels were similar at both effector:target ratios. In contrast, a weak infiltration of HD NKs was observed in NIB140 spheroids and slightly higher in the NCH421k spheroids (Fig. 4c). Although not reaching statistical significance, IL-2 pre-activation slightly improved the infiltration of HD NKs. The extent of NK-92 and HD NK cell infiltration was also confirmed by immunofluorescent detection of CD45-positive NK cells in FFPE sections of spheroid-NK cell co-cultures (Fig. 4d-e). Overall, most NK cells were concentrated near the spheroid surface, but some NK cells were also observed deep within the spheroids, which indicates their ability to penetrate the cell-dense GB spheroid structure. We also showed that infiltrated NK-92 cells proliferated in the GB spheroids (Fig. S5). To further compare the immune-attracting potential of NCH421k and NIB140 spheroids, we also set their direct co-cultures with GB patient PBMCs (Fig. S6). Similar to NK-92 cells, IL-2-activated PBMCs infiltrated at much higher levels into NIB140 spheroids, compared to NCH421k spheroids.

**Figure 4.**
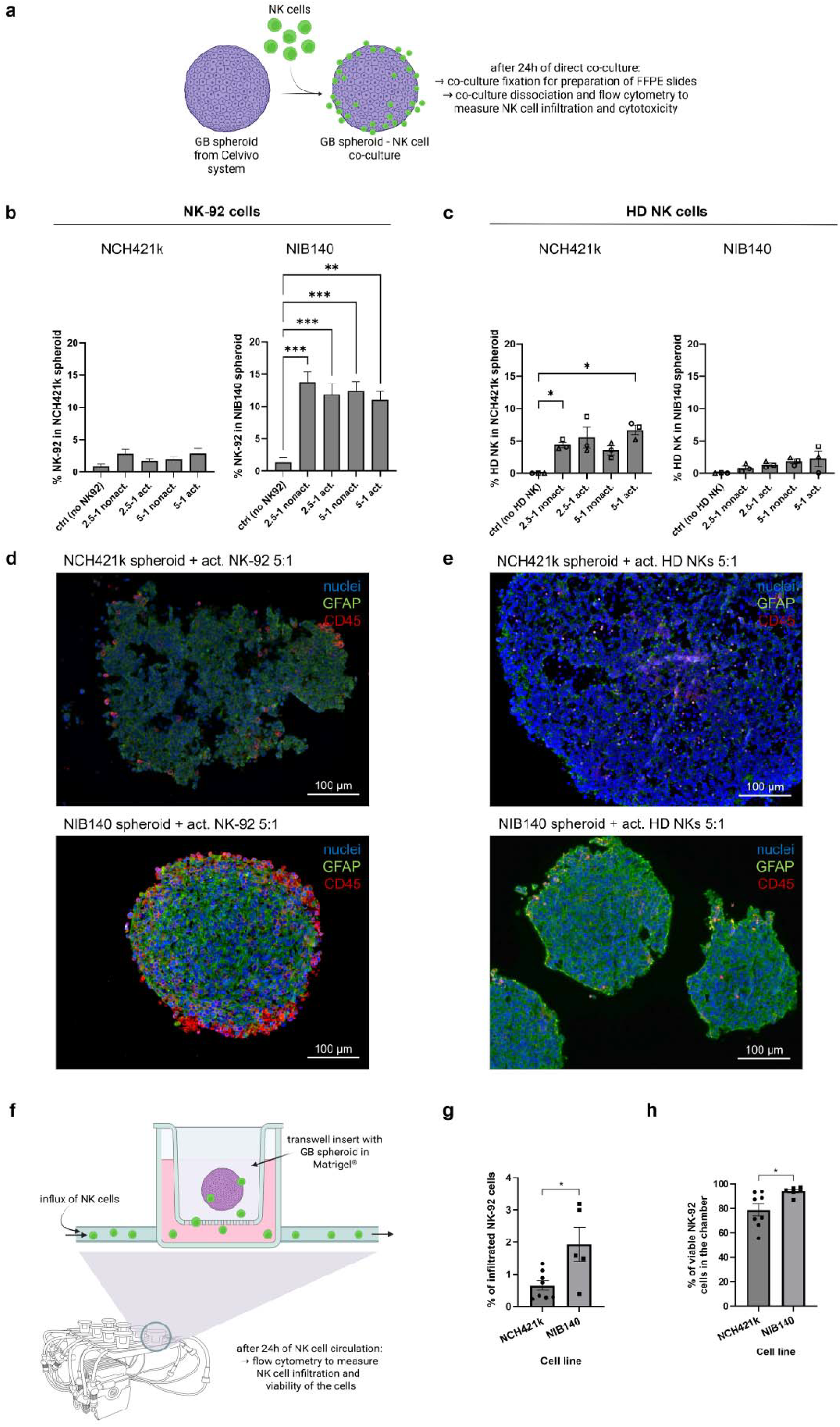
NK cell infiltration in NCH421k and NIB140 spheroids. **a)** Schematic representation of the direct GB spheroid-NK cell co-culture set-up. Created with bioRender.com. **b)** NK-92 cell infiltration into NCH421k and NIB140 spheroids as determined by flow cytometry after the 24 h co-culture. Data are presented as means ±⍰SEM of 6 biological replicates. Ordinary one-way ANOVA was used for statistical analysis. **c)** HD NK cell infiltration into NCH421k and NIB140 spheroids as determined by flow cytometry after the 24 h co-culture. Data are presented as means ±⍰SEM of 3 biological replicates. Each data point and its shape correspond to HD NK cells from an individual donor. Ordinary one-way ANOVA was used for statistical analysis. **d)** Immunofluorescent staining of CD45-positive NK-92 cells on FFPE slides of GB spheroid-NK-92 co-cultures. **e)** Immunofluorescent staining of CD45-positive HD NK cells on FFPE slides of GB spheroid-HD NK cell co-cultures. **f)** Schematic representation of the organ-on-chip platform mimicking the blood flow and influx of NK-92 cells into the tumour. Created with bioRender.com. **g)** Infiltration of NK-92 cells into chambers containing NCH421k/NIB140 spheroids. After 24 h of NK-92 circulation, the percentage of NK-92 was determined by flow cytometry. Data are shown as mean ±⍰SEM and data points represent each of the 5–8 biological replicates. Unpaired t-test was used for statistical analysis. **h)** Viability of NK-92 cells infiltrated into the chambers containing NCH421k/NIB140 spheroids. Data are shown as mean ±⍰SEM and data points represent each of the 6–8 biological replicates.

The ability of NK-92 cells to infiltrate GB spheroids was additionally tested in a dynamic organ-on-chip platform mimicking the blood flow and influx of immune cells from blood circulation into the tumour (Fig. 4f-h). NCH421k or NIB140 spheroids were embedded in Matrigel in the porous transwell insert and NK-92 cells were applied to the circulation system (Fig. 4f). After 24 h of constant NK-92 cell circulation, flow cytometry was used to quantify NK-92 cells that infiltrated into the transwell insert and GB spheroids. In line with the results in direct co-cultures, also in the setting of this physiologically relevant *in vitro* model, NIB140 spheroids exhibited a higher attraction for NK-92 cells compared to NCH421k spheroids (Fig. 4g). Importantly, the infiltration of NK-92 cells was measured at the same concentration of FBS in the chambers for both spheroid types, as FBS *per se* significantly increased infiltration (Fig. S7a). The percentage of dead GB cells in the insert was slightly higher in NCH421k spheroids compared to NIB140 spheroids (Fig. S7b). Interestingly, infiltrated NK cells were less viable in chambers containing NCH421k spheroids compared to chambers with NIB140 spheroids, but viability was still approximately 80% or higher (Fig. 4h).

### 3.4 NK cells show differences in cytotoxicity against GB spheroids derived from different cell lines

In addition to measuring the ability of NK cells to infiltrate GB spheroids, we also measured their cytotoxicity after 24 h of direct co-culture (Fig. 5). In contrast to the observed spheroid infiltration levels, NK-92 cells exhibited much higher cytotoxicity against NCH421k spheroids compared to NIB140 spheroids (Fig. 5a). NK-92 cytotoxicity against NCH421k spheroids increased with increasing E:T ratio. IL-2-activated NK-92 cells at the 5:1 E:T ratio killed over 30% of NCH421k cells in spheroids. In the case of NIB140 spheroids, no significant increase in the proportion of dead cells was observed in the NK-92 co-cultures compared to control, indicating a profound resistance of NIB140 cells to NK-92 cell-mediated lysis. In terms of HD NK cell cytotoxicity, there were substantial differences among the donors (Fig. 5b). Overall, similar to the NK-92 cells, also the HD NKs were more cytotoxic against NCH421k spheroids compared to spheroids from NIB140 cells, which were highly resistant to HD NK cell-mediated killing. The cytotoxic activity of both NK-92 cells and HD NKs against NCH421k was slightly increased by IL-2 pre-activation. Consistent with higher cytotoxicity of NK-92 and HD NKs against NCH421k spheroids compared to NIB140 spheroids, we also detected an increase of IFN-γ in the media of NCH421k co-cultures, while its levels were unchanged or even decreased in NIB140 co-cultures (Fig. S8).

**Figure 5.**
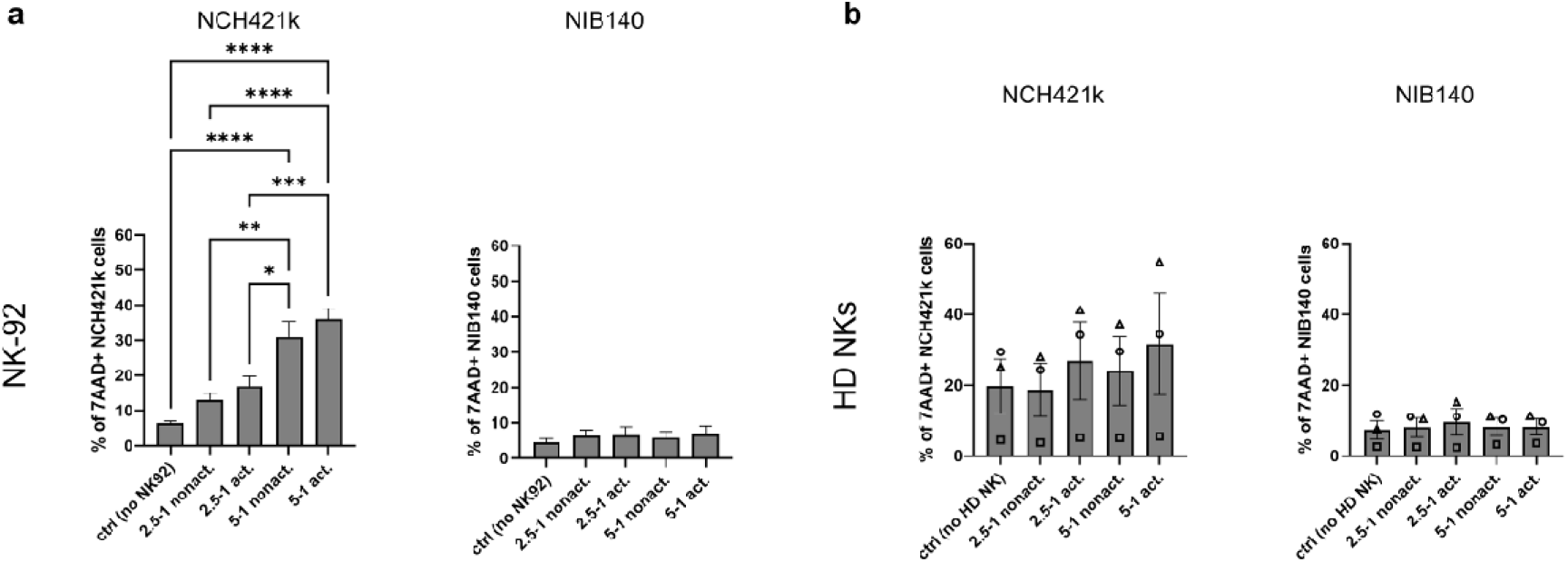
NK cell cytotoxicity against NCH421k and NIB140 spheroids. **a)** Flow cytometric analysis of NK-92 cytotoxicity against NCH421k and NIB140 spheroids. Data are presented as mean ±⍰SEM of 6 biological replicates. Ordinary one-way ANOVA was used for statistical analysis. **b)** Flow cytometric analysis of HD NK cells cytotoxicity against NCH421k and NIB140 spheroids. Data are presented as means ±⍰SEM of 3 biological replicates. Each data point and its shape correspond to HD NK cells from an individual donor. Ordinary one-way ANOVA was used for statistical analysis.

### 3.5 GB spheroids derived from different cell lines secrete variable levels of soluble factors that affect NK cell infiltration and cytotoxicity

Since the infiltration and cytotoxic activity of NK cells may be influenced by soluble factors released from cancer cells, we tested whether conditioned media from GB cell lines affects NK cell cytotoxicity. Calcein release assay was performed with NK-92 cells and HD NKs pre-cultured in GB cell conditioned media for 48h prior to the onset of the assay. K562 cells were used as target cells. NIB140- and NCH421k-conditioned media significantly decreased cytotoxicity of NK-92 cells by over or nearly 50%, respectively (Fig. 6a). In parallel, lower levels of activating receptor NKp30 were detected on NK-92 cells cultured in GB cell-conditioned media (Fig. 6b). NIB140 (but not NCH421k) conditioned medium also increased NKG2D and decreased PD-1 on NK-92 cells (Fig. S9). Conditioned medium from either of the cell lines did not affect the cytotoxicity of HD NKs (Fig. S10), probably due to high activation of HD NK cells.

**Figure 6.**
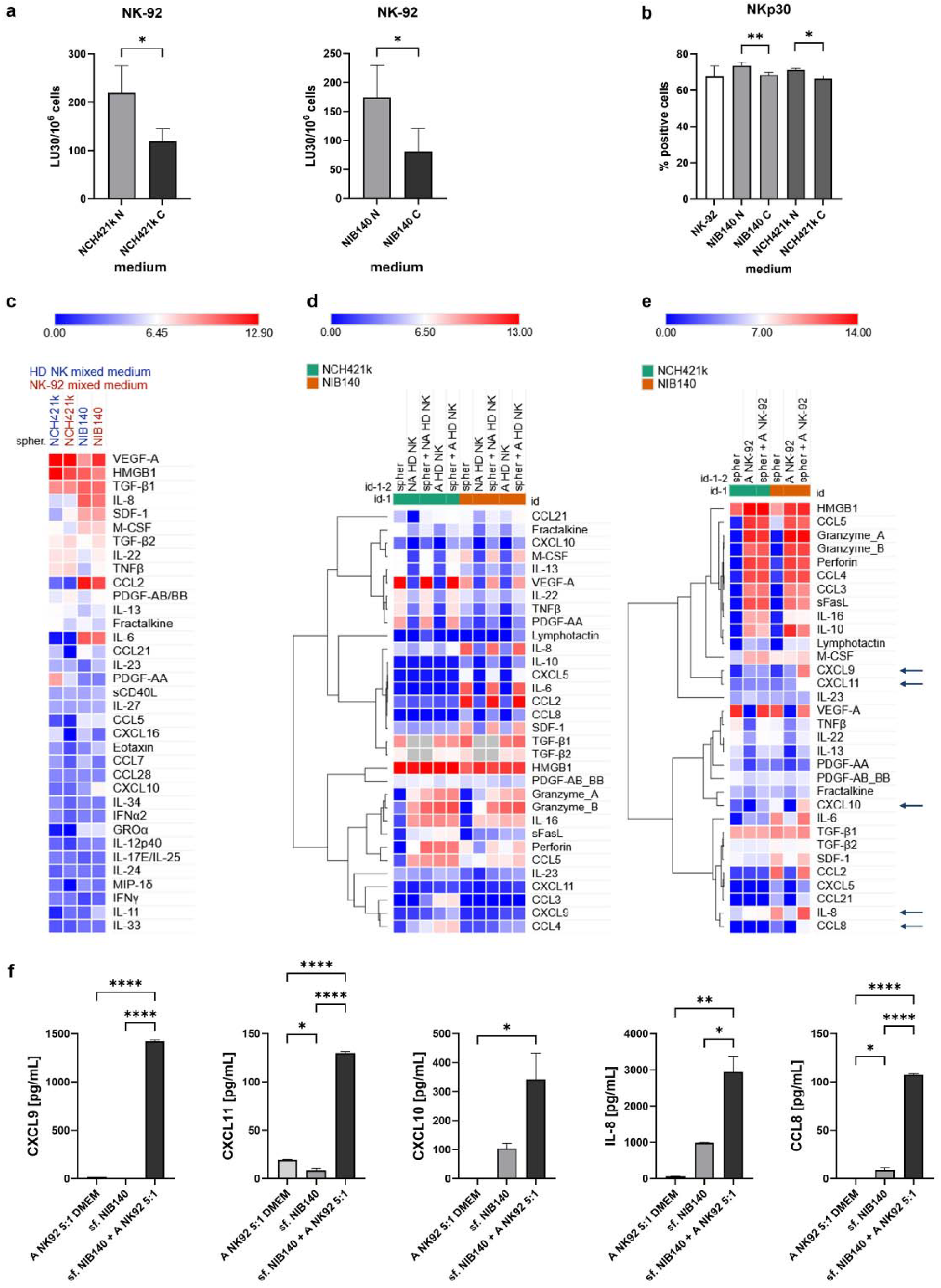
Soluble factors in GB spheroid cultures and their 24 h co-cultures with NK cells. **a)** The effect of NCH421k- and NIB140-conditioned medium on cytotoxicity of NK-92 cells against K562 cells. Data are presented as means ±⍰SEM of 3–4 biological replicates. Paired t-test was used for statistical analysis. N – nonconditioned medium, C – conditioned medium. **b)** Expression of NKp30 on NK-92 cells cultured in NCH421k- or NIB140-conditioned medium. Data are presented as mean ±⍰SEM of 3– 4 biological replicates. Paired t-test was used for statistical analysis of differences in expression between nonconditioned and conditioned media. N – nonconditioned medium, C – conditioned medium. **c)** Analysis of cytokine secretion in NCH421k and NIB140 spheroids. log2(x+1)-transformed cytokine concentrations (averaged from two technical replicates) are presented in the heatmap. **d)** Analysis of secreted cytokines in the 24 h spheroid-HD NK cell co-cultures. Only cytokines with concentration ≥ 50 pg/mL in at least one of the conditions were included. log2(x+1)-transformed cytokine concentrations (averaged from two technical replicates) are presented in the heatmap. Cytokines were hierarchically clustered based on the “one minus Pearson correlation” metric in Morpheus. **e)** Analysis of secreted cytokines in the 24 h spheroid-NK-92 cell co-cultures. Only cytokines with concentration ≥ 50 pg/mL in at least one of the conditions were included. log2(x+1)-transformed cytokine concentrations (averaged from two technical replicates) are presented in the heatmap. Cytokines were hierarchically clustered based on the “one minus Pearson correlation” metric in Morpheus. **f)** CXCL9, CXCL11, CXCL10, IL-8 and CCL8 were prominently upregulated in co-cultures of NK-92 cells with NIB140 spheroids. Data are presented as mean ±⍰SEM of two technical replicates. Ordinary one-way ANOVA was used for statistical analysis.

To further inspect the possibly immunomodulating factors released from GB cells, we analysed the concentrations of cytokines, chemokines and cytotoxicity-related factors in media supernatants of GB spheroids and their 24 h co-cultures with NK-92 and HD NK cells (Fig. 6c-f, Fig. S11-13). Spheroids derived from NCH421k (i.e., GB stem-like) and NIB140 (i.e., differentiated) GB cells varied significantly in their cytokine secretion profiles (Fig. 6c). While spheroids from both cell lines secreted high levels of vascular endothelial growth factor A (VEGF-A), high mobility group protein B1 (HMGB1) and transforming growth factor beta (TGF-β), NIB140 spheroids secreted much more pro-inflammatory cytokines such as C-C motif chemokine ligand 2 (CCL2), IL-6, IL-8, stromal cell-derived factor 1 (SDF-1), and macrophage colony-stimulating factor (M-CSF) compared to NCH421k spheroids.

The secretion of several factors was modulated in the media of GB spheroid-NK cell co-cultures compared to mono-culture controls (Fig. 6d-e, Fig. S11, Fig. S12). In both NK-92 and HD NK cell cocultures with GB spheroids, cytotoxicity-related factors including granzymes A and B, perforin, and soluble Fas ligand (sFasL) clustered with IL-16 and CCL5 (Fig. 6d-e). Additionally, CCL3, CCL4, IL-10, and lymphotactin were within the cluster in NK-92 co-cultures. IL-8 and M-CSF were significantly upregulated in all co-cultures, except for NIB140 spheroids with HD NKs (Fig. S12). Downregulation of tumour necrosis factor beta (TNF-β), IL-10, IL-13, perforin, SDF-1, sFasL and VEGF-A, and upregulation of C-X-C motif chemokine ligand 11 (CXCL11) were specifically observed in NK-92 co-cultures, but not HD NK co-cultures. Platelet-derived growth factor (PDGF) was significantly downregulated only in NK-92 co-culture with NCH421k. Overall, the most striking upregulations were observed for IL-8, CXCL9, CXCL11, CCL8, and CXCL10 in NK-92 cell co-cultures with NIB140 spheroids (Fig. 6f, Fig. S13).

### 3.6 GB patient NK cells express less activating NK cell receptors and TIGIT compared to NK cells from healthy donors

As the function of NK cells also significantly depends on the presence of activating and inhibitory receptors on their surface, we analysed the expression of selected NK receptors on IL-2 activated NK-92 cells, HD NK cells and, in addition, GB patient NK cells (Fig. 7). The expression of receptors varied between the three NK cell sources (Fig. 7a, Fig. S14a). NK-92 cells expressed activating receptors NKp30 and NKG2D, but also inhibitory (co)receptors CD94, CD96 and TIGIT. A majority of HD NKs expressed Fc receptor CD16; varying levels of activating receptor DNAM-1 and inhibitory receptors CD94, TIGIT, and PD-1 were also detected. Compared to HD NKs, GB patient NKs expressed less CD16, NKp30, TIGIT, NKG2D, and DNAM-1, and more immune check-point PD-1 (Fig. 7a, Fig. S14b). Expectedly, IL-2 activation of HD NKs significantly increased the expression of activating receptors NKG2D and NKp30 and decreased the expression of PD-1 (Fig. 7b, Fig. S14c).

**Figure 7.**
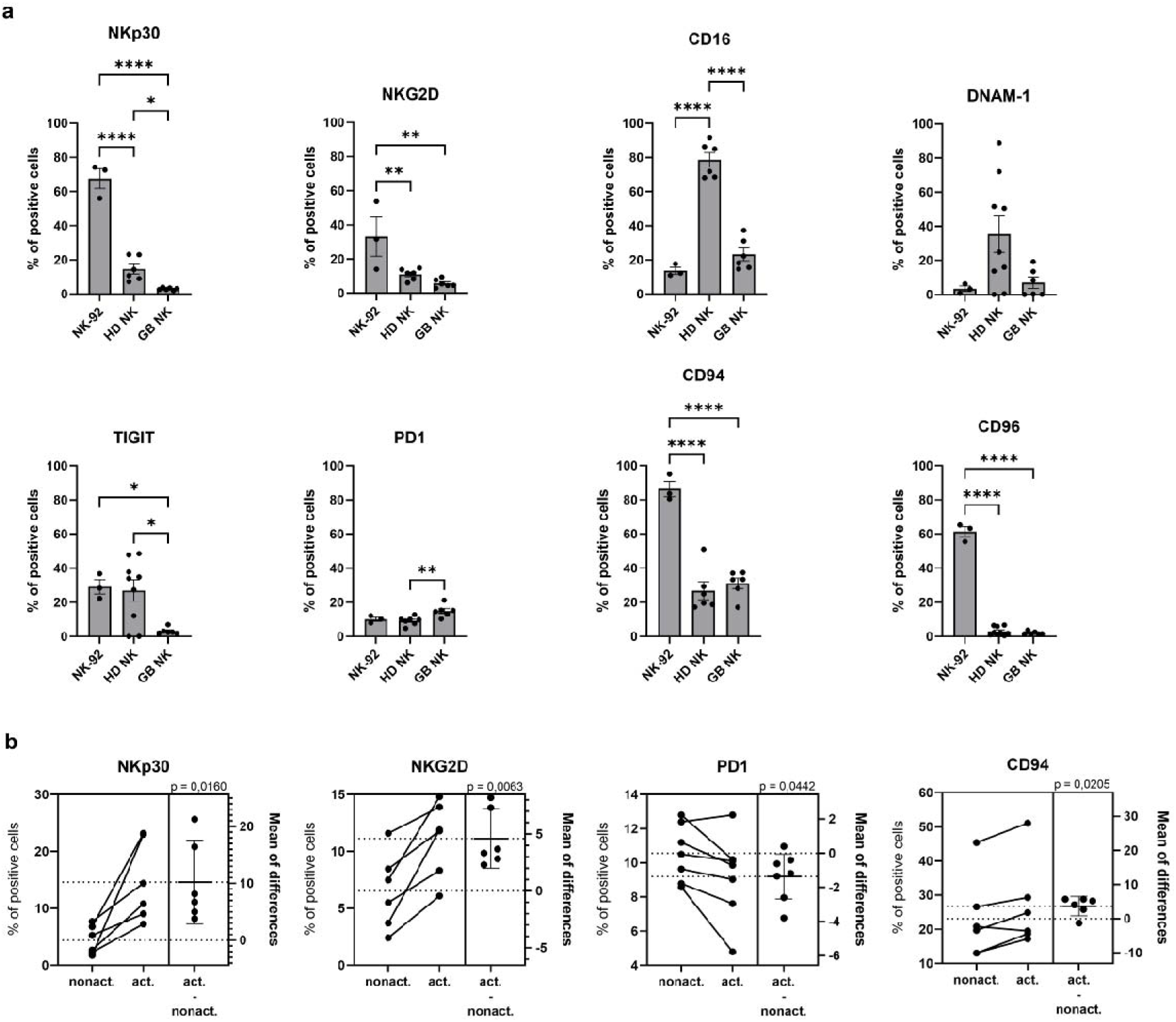
Expression of NK cell receptors on IL-2-activated NK cells from different sources. **a)** Comparison of receptor ex pression on IL-2-activated NK-92 cells (N=3), HD NK cells (N=6–9) and NK cells from GB patients (N=6). Data are presented as mean ±⍰SEM and dots represent biological replicates. Ordinary one-way ANOVA was used for statistical analysis. **b)** Estimation plots showing the expression of NK cell receptors on HD NK cells, affected by IL-2-activation (N=6).

### 3.7 The CD155-TIGIT/DNAM-1 axis is an important mediator of HD NKs’ cytotoxicity against NCH421k cells

As high CD155 expression was observed in standard and spheroid cultures of GB cells and since decreased expression of CD155 receptors TIGIT and DNAM-1 was detected on NK cells from GB patients, we further explored the role of the CD155 axis in regulation of NK cell cytotoxicity against GB cells. According to the TCGA and CGGA data, *PVR* (the gene that encodes CD155) is upregulated in GB compared to low-grade gliomas (Fig. 8a, Fig. S15a) and high levels of *PVR* correlate with poor prognosis of glioma patients (Fig. 8b, Fig. S15b).

**Figure 8.**
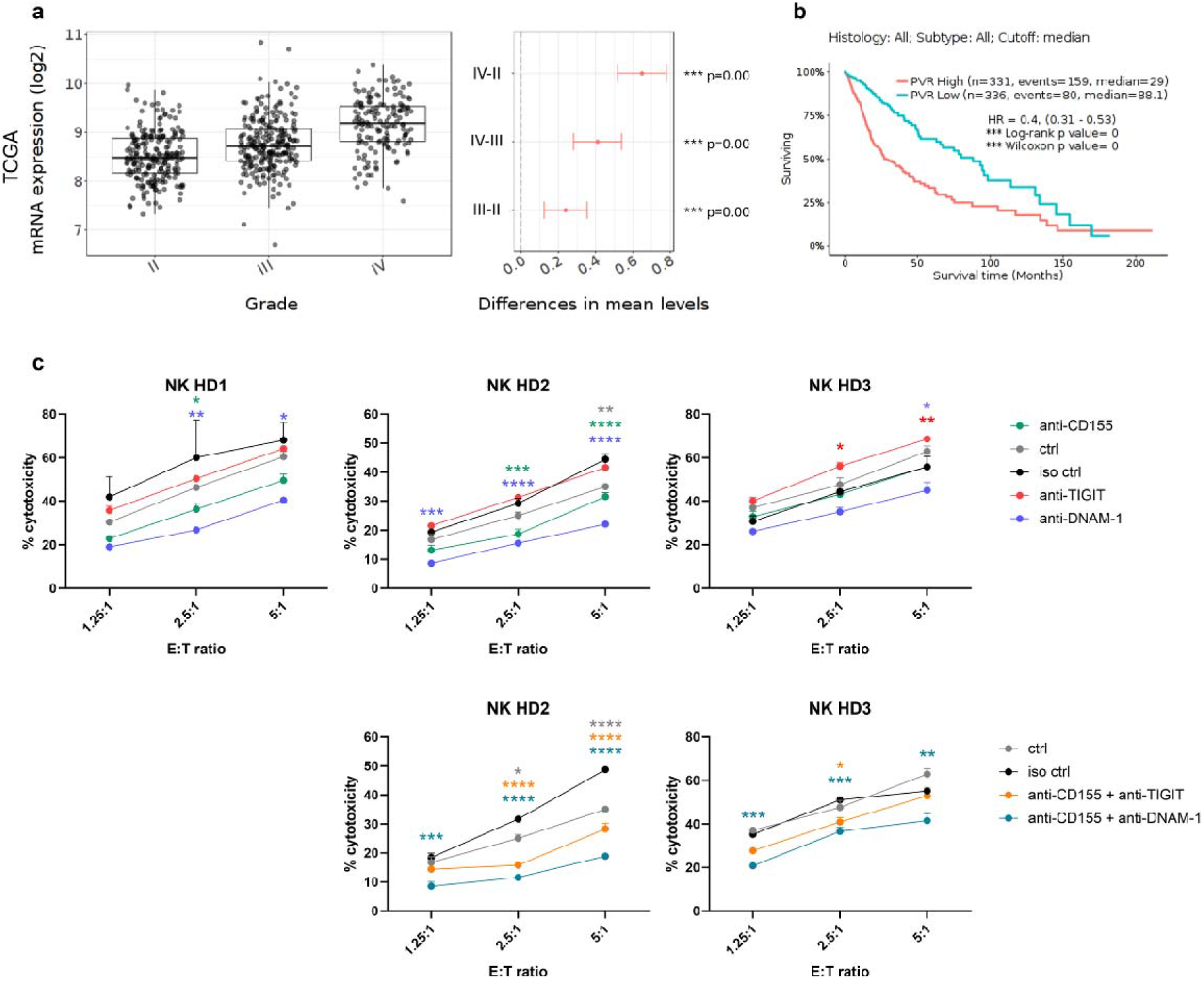
The CD155 axis regulates the cytotoxicity of HD NK cells against NCH421k cells. **a)** According to the TCGA data, *PVR* is upregulated in GB compared to lower-grade gliomas. Tukey’s Honest Significant Difference were calculated in GlioVis. **b)** Higher expression of *PVR* is associated with shorter survival of glioma patients. Kaplan-Meier estimator survival analysis was performed in GlioVis. **c)** Results of calcein release assay performed with NK cells from 3 different healthy donors in which CD155, TIGIT, and DNAM-1 or their combinations were blocked by antibodies and NCH421k cells were used as target cells. Data are shown as mean ±⍰SEM of three technical replicates. 2-way ANOVA with Tukey’s multiple comparisons test was used for statistical analysis.

We used calcein release assay to test the effects of antibody-based blockade of CD155 on NIB140/NCH421k cells and its receptors TIGIT or DNAM-1 on HD NK cells (Fig. 8c, Fig. S16). The assay was performed with NKs from three healthy donors, which expressed varying levels of TIGIT and DNAM-1 but did not express CD96 (Fig. S17). Blocking components of the CD155 axis had a much greater effect on cytotoxicity against NCH421k cells (Fig. 8c) compared to NIB140 cells (Fig. S16). Blocking CD155 decreased cytotoxicity against NCH421k cells in NKs from two of three donors (Fig. 8c). Blockade of DNAM-1 consistently decreased HD NK cell cytotoxicity against NCH421k cells in NKs from all three donors, while blocking TIGIT upregulated NK cell activity only in the case of NKs from healthy donor three, which expressed the highest level of TIGIT among the donors. Simultaneous blockade of CD155 and TIGIT or, to a greater extent, CD155 and DNAM-1, severely impaired the ability of HD NKs to eliminate NCH421k cells.

## 4. Discussion

In this study, we established a protocol for reproducible production and dynamic culture of uniformly sized GB spheroids in the Celvivo Clinostar system. Uniform spheroid size is particularly important for reliability and reproducibility of further studies, such as assessing the responses of spheroids to therapies, in which their variable size may significantly distort experimental results (28). Another key advantage of our approach is that the spheroids in the Celvivo Clinostar system can be cultured for substantially longer time periods compared to other protocols in which spheroids are maintained under static conditions. Thus, our method is well suited for specific long-term experiments and studies of chronic treatments. In the present study, spheroids were produced from several distinct cell lines, including suspension-cultured as well as adherent GB cell lines, implying a broad applicability of the approach. In time, the architecture of our spheroid models became increasingly heterogeneous. Like GB tumours *in vivo*, our models developed necrotic and hypoxic regions and also (at least partly) recapitulated the heterogeneity of cancer cell states. Specifically, distinct zones of highly and poorly proliferative cells were observed and cells expressed varying levels of stemness marker SOX2.

Cancer cell spheroids reproduce the 3D conformation of the tumour, including cell-cell interactions, extracellular matrix, and gradients of nutrients and oxygen. As immune cell function is severely affected by nutrient and oxygen deprivation and such conditions are common in tumours due to their extensive growth and abnormal vasculature (29–32), using spheroid models in immunotherapy research may reveal insights into immune cell responses under conditions close to those observed *in vivo*. Importantly, also the expression of immune-related molecules on cancer cells can be affected by the culture type (2D vs. 3D) (33). It has previously been reported that GB cells cultured in 3D upregulate the expression of immunosuppressive molecules, such as HLA-E and MICA/B (34,35). In line, we observed consistent downregulation of ligands for several activating NK cell receptors (CD155, B7-H6 and ULBPs) in GB spheroid cultures. Interestingly, major differences were also observed in the NCH421k cell line, which is classically cultured in the form of small multicellular aggregates, implying that not just inter-cellular connections, but also spheroid size is an important factor influencing the cancer cell immune phenotype. Compared to the standard adherent NIB140 culture, cells in NIB140 spheroids expressed significantly lower levels of HLA-A/B/C. Downregulation or loss of HLA class I molecules is one of the main mechanisms of tumour immune evasion, which hides cancer cells from recognition by T cells, but may simultaneously increase their sensitivity to NK cells (36).

Based on the differences in expression of stemness related genes, NCH421k and NIB140 spheroids were used as proxies of GB stem-like and differentiated GB cells, respectively. We showed that NIB140 spheroids (i.e., differentiated GB cells) produced much higher levels of pro-inflammatory and immune-attracting factors (CCL2, IL-8, IL-6, SDF-1, M-CSF) compared to the stem-like NCH421k spheroids. As also indicated by others (37), our results strongly suggest that differentiated GB cells should be considered in development of novel immunotherapeutic approaches. Although GB stem cells are generally viewed as the most critical therapeutic targets, differentiated GB cells may significantly affect the immune landscape within the TME, shape adaptive immune responses and thus also influence the success of immunotherapies. For example, CCL2 may attract several effectors of immunosuppression, including macrophages, regulatory T cells (Tregs), and myeloid-derived suppressor cells (MDSCs), (38–40). Similarly, MDSCs can also be recruited by IL-8 (41,42) and their accumulation and immune inhibitory functions are enhanced by IL-6 (43).

In the 24 h direct co-culture of GB spheroids and NK cells, we observed substantial differences in NK-92 and HD NK cell infiltration and cytotoxic capacities. While the infiltration of HD NKs correlated with their cytotoxicity, no such trend was observed with NK-92 cells. Even though NK-92 cells profoundly killed NCH421k cells in the spheroids, few NK-92 cells were infiltrated within the spheroids. Conversely, NK-92 cells successfully infiltrated into the dense structure of NIB140 spheroids, which were highly resistant against NK-92 mediated killing. These results suggest that the cytotoxic activity does not necessarily correlate with the ability of NK cells to penetrate the spheroid and that measuring NK cell infiltration and their cytotoxicity should be considered as two distinct read-outs.

The remarkable infiltration of NK-92 cells into NIB140 spheroids compared to NCH421k spheroids may be attributed to a number of potentially NK cell attracting cytokines secreted at high levels in NIB140 spheroid cultures. These include, for example, CCL2 (44), IL-8 (45,46), and SDF-1 (47). Additionally, in NK-92 co-cultures with NIB140 spheroids there was significant increase in levels of IL-8, CCL8 and IFN-γ-inducible cytokines CXCL9, CXCL11 and CXCL10. CXCL9, CXCL10 and CXCL11 may further potentiate NK cell infiltration as they bind to CXCR3, a receptor with a fundamental role in NK cell trafficking (48). Speculating that the CXCR3 ligands are also upregulated upon interaction of GB and NK cells in GB TME *in vivo*, they could also attract activated T cells and other leukocytic subtypes which are known to express CXCR3. Of note, CXCR3 is also expressed on GB cells and its isoforms may have either pro- or anti-tumorigenic functions (49).

The cytotoxic potential of both NK-92 and HD NK cells was higher against NCH421k spheroids compared to NIB140 spheroids. This is in line with a number of studies which demonstrate that NK cells can recognize and kill GB stem-like cells *in vitro* and that GB stem-like cells are more susceptible to NK cells compared to their differentiated counterparts (19,50,51). We have shown that GB stem-like cells upon differentiation with serum and IFN-γ become more resistant to NK-cell mediated killing (19). Interestingly, despite a profound resistance of NIB140 spheroids against HD NKs and NK-92 cells, both NK cell sources showed dose-dependent cytotoxicity against single NIB140 cells dissociated from spheroids in calcein release assay. This may imply that the activity of NK-92 cells is specifically obstructed within the 3D environment of NIB140 cells and highlights the importance of the 3D research models. *In vivo*, the immune phenotype of GB-infiltrating NK cells is distinct to NK cells from matched peripheral blood. Within the tumour, NK cells downregulate activating receptors and reduce IFN-γ secretion (52–54), which may also occur upon infiltration into GB spheroids *in vitro*. In the spheroids, NK-92 cell activity may be suppressed by either 1) physical interactions within the cell-dense structure of the spheroid or/and 2) by NK-suppressive factors released by GB cells that demonstrate a greater effect in the 24 h co-culture experiment compared to the 3 h calcein release assay. Indeed, NIB140 spheroids (but not NCH421k spheroids) secreted high levels of IL-6 and IL-8, which are known to impair NK cell activity (55). Compared to NCH421k spheroids, NIB140 spheroids also secreted more TGF-β1, one of the major suppressors of NK cell function in GB (24,25). Culturing NK-92 cells in NIB140 (as well as NCH421k) conditioned medium inhibited their cytotoxicity against K562 cells, indicating a profound effect of GB soluble factors in regulation of NK-92 cell activity. Interestingly, neither NIB140 nor NCH421k conditioned medium affected the cytotoxicity of HD NKs, implying that (compared to NK-92 cells) these cells are in high activation state and may be less prone to inactivation by GB soluble factors.

The repertoire of activating and inhibitory receptors on the NK cell surface is an important factor that largely affects NK cell function. NK cell receptors may be dysregulated on NK cells in cancer patients (56,57). Inspecting NK cells from GB patients revealed that, compared to HD NKs, these cells expressed less CD16, NKp30, TIGIT, NKG2D, and DNAM-1 and more PD-1. Downregulation of activating receptors on GB patient NKs was described previously (23) and we previously reported reduced cytotoxicity and decreased IFN-γ secretion in GB patient NKs when co-cultured with cancer cells (19). However, our attention was caught by reduced levels of TIGIT and DNAM-1, which both bind CD155 – a ligand that was highly expressed in standard as well as spheroid GB cultures. We thus decided to further inspect the role of the CD155 axis in NK cell cytotoxicity against GB cells.

CD155 (also known as the poliovirus receptor or “PVR”) is a type I transmembrane glycoprotein from the nectin and nectin-like protein family. Its expression is upregulated in many cancers which often correlates with poor patient prognosis (58,59). In GB, CD155 has been associated with a number of diverse functions such as cancer cell adhesion, migration, invasion, and proliferation (60–62). Importantly, CD155 also regulates immune responses of NK cells and other lymphocytes. It can bind to several receptors, including the inhibitory TIGIT, the activating DNAM-1, and CD96, a receptor with dual roles (58,59). Inspecting the role of the CD155 axis in regulation of NK cell cytotoxicity against GB cells showed that although the expression of the ligand was higher on NIB140 cells compared to NCH421k cells, blocking CD155 or its receptors only showed convincing effects on NK cell cytotoxicity against NCH421k cells.

The inhibitory receptor TIGIT generally has a stronger affinity for CD155 compared with the activating receptor DNAM-1 (63). Although TIGIT and, to a greater extent, DNAM-1 were both expressed on NK cells from all three healthy donors, a decrease in cytotoxicity upon blockade of CD155 on NCH421k cells suggests that in net effect, the ligand predominantly induced activating signalling. Blocking DNAM-1 significantly decreased HD NK cytotoxicity against NCH421k cells, with an effect stronger than that observed with blocking CD155 itself. This implies that DNAM-1 probably also activates NK cells by binding other ligands, e.g., CD112, which we also detected on NCH421k cells. Blocking TIGIT only moderately increased cytotoxicity of NKs from healthy donor three, which expressed the highest level of TIGIT (and DNAM-1) among the three donors, while it showed no effect in NKs from the other two donors. Our observations are in line with a recent study which concluded that TIGIT *per se* does not directly inhibit NK cells upon interaction with CD155 on GB cells, but rather acts as a decoy receptor, preventing the interaction with DNAM-1 (64). Thus, blocking TIGIT alone may not reliably restore NK activity against GB. All in all, our results may imply that the CD155 axis is an important regulator of NK cell cytotoxicity against GB stem-like cells, although additional cell lines should be tested for confirmation of our findings. Nevertheless, we can conclude that the importance of the axis depends on the overall immune phenotype of cancer cells consisting of a diverse repertoire of additional ligands for NK cell receptors.

## 5. Conclusions

In the present study, we established a general protocol for reproducible production and dynamic culture of uniformly sized GB spheroids in the Celvivo Clinostar system. The model was used to explore the crosstalk between GB and NK cells. Several ligands for NK cell receptors were differentially expressed in spheroids compared to cells in standard culture, affecting their interactions with NK cells. Although NK cells infiltrated GB spheroids, their infiltration capacity did not necessarily correlate with their ability to eliminate GB cells. GB cell lines secreted variable levels of soluble factors that impaired NK cell cytotoxicity. Spheroids derived from differentiated GB cells secreted higher levels of immune-attracting, pro-inflammatory and NK cell-inhibiting cytokines compared to GB stem-like spheroids, indicating a profound ability of differentiated GB cells to shape the GB immune milieu. Lastly, the CD155-TIGIT/DNAM-1 axis was found to be an important mediator of NK cell cytotoxicity against GB stem-like cells. Collectively, our results highlight important factors in the GB-NK cell communication and present a groundwork for further targeted research.

## Abbreviations

ADCC: antibody dependent cellular cytotoxicity
B7-H6: B7 homolog 6
BSA: bovine serum albumin
CA9: carbonic anhydrase 9
CCL: C-C motif chemokine ligand
CD112: nectin-2
CD155: poliovirus receptor
CD16 (FcγRIII): cluster of differentiation 16 (Fc gamma receptor III)
CD44: CD44 protein
CD54 (ICAM-1): cluster of differentiation 54 (intercellular adhesion molecule 1)
CD96 (TACTILE): cluster of differentiation 96 (T-cell surface protein tactile)
CXCL: C-X-C motif chemokine ligand
DNAM-1: DNAX Accessory Molecule-1
EDTA: ethylenediaminetetraacetic acid
FBS: foetal bovine serum
FFPE: formalin fixed paraffin embedded
GB: glioblastoma
GFAP: glial fibrillary acidic protein
HD: healthy donor
HIF1α: hypoxia inducible factor 1 subunit alpha
HLA: human leukocyte antigen
HMGB1: high mobility group protein
B1 IFN-γ: interferon-gamma
IL-2: interleukin 2
Ki-67: proliferation marker protein Ki-67
KIR: killer cell immunoglobulin-like receptor
M-CSF: macrophage colony-stimulating factor
MICA: MHC class I polypeptide-related sequence A
MICB: MHC class I polypeptide-related sequence B
NHA: normal human astrocytes
NK: natural killer
NKG2D: NKG2-D type II integral membrane protein
NKp30 (NCR3): natural cytotoxicity triggering receptor 3
OLIG2: oligodendrocyte transcription factor 2
PBMCs: peripheral blood mononuclear cells
PBS: phosphate-buffered saline
PDGF: platelet-derived growth factor
PD-L1: programmed cell death 1 ligand 1
SDF-1: stromal cell-derived factor 1
SEM: standard error of the mean
sFasL: soluble Fas ligand
SOX2: SRY-box transcription factor 2
TGF-β: transforming growth factor beta
TIGIT: T cell immunoreceptor with Ig and ITIM domains
TME: tumour microenvironment
TNF-α: tumour necrosis factor-α
TNF-β: tumour necrosis factor beta
ULBP: UL16 binding protein
VEGF-A: vascular endothelial growth factor A

## Declarations

### Ethics approval and consent to participate

The study was approved by the National Medical Ethics Committee of the Republic of Slovenia (approval No. 0120-190/2018-2711-41). Experiments with human buffy coats were performed in accordance with the approval no. 0120-279/2017-3.

### Consent for publication

Not applicable.

### Availability of data and materials

No datasets were generated or analysed during the current study.

### Competing interests

The authors declare no competing interests.

### Funding

This study was supported by the Slovenian Research and Innovation Agency (programme and research grants P1-0245, J3-4504, NC-25002, N3-0394, and young researcher grant to AH), European Union’s projects CutCancer 101079113, SPACETIME 101136552, UNCAN-CONNECT 101215206, GenomeMET 22HLT06.

### Authors’ contributions

A.H., M.N, and B.B. conceptualised the study. A.H., T.K.M., P.Ž., B.M., Š.K., E.S., and M.P.N. designed and performed experiments and analysed the obtained data. U.Š., A.P., and B.P. provided PBMCs/blood samples from healthy donors and GB patients and performed experiments. M.N and B.B. supervised the study. A.H., M.N., and B.B. wrote the first draft of the manuscript. All authors reviewed, edited and approved the submitted manuscript.

## Acknowledgements

We acknowledge the support of COST actions IMMUNO-model CA21135 and Net4Brain CA22103.

## Notes

### Competing Interest Statement

The authors have declared no competing interest.

## References

1. van den Bent MJ, Geurts M, French PJ, Smits M, Capper D, Bromberg JEC, et al. Primary brain tumours in adults. Lancet. 2023;402(10412):1564–79.

2. Miller KD, Ostrom QT, Kruchko C, Patil N, Tihan T, Cioffi G, et al. Brain and other central nervous system tumor statistics, 2021. CA Cancer J Clin. 2021;71(5):381–406.

3. Tang L, Huang Z, Mei H, Hu Y. Immunotherapy in hematologic malignancies: achievements, challenges and future prospects. Signal Transduct Target Ther. 2023;8(1):1–39.

4. Guha P, Heatherton KR, O’connell KP, Alexander IS, Katz SC. Assessing the Future of Solid Tumor Immunotherapy. Biomedicines. 2022;10(3):655.

5. Liu Y, Zhou F, Ali H, Lathia JD, Chen P. Immunotherapy for glioblastoma: current state, challenges, and future perspectives. Cell Mol Immunol. 2024;21(12):1354–75.

6. Akter F, Simon B, de Boer NL, Redjal N, Wakimoto H, Shah K. Pre-clinical tumor models of primary brain tumors: Challenges and Opportunities. Biochim Biophys acta Rev cancer. 2020;1875(1):188458.

7. Majc B, Novak M, Jerala NK, Jewett A, Breznik B. Immunotherapy of Glioblastoma: Current Strategies and Challenges in Tumor Model Development. Cells. 2021;10(2):265.

8. Wang Q, Yuan F, Zuo X, Li M. Breakthroughs and challenges of organoid models for assessing cancer immunotherapy: a cutting-edge tool for advancing personalised treatments. Cell Death Discov. 2025;11(1):1–13.

9. Wouters R, Bevers S, Riva M, De Smet F, Coosemans A. Immunocompetent Mouse Models in the Search for Effective Immunotherapy in Glioblastoma. Cancers. 2020;13(1):19.

10. Paterson K, Zanivan S, Glasspool R, Coffelt SB, Zagnoni M. Microfluidic technologies for immunotherapy studies on solid tumours. Lab Chip. 2021;21(12):2306–2329.

11. Wolf NK, Kissiov DU, Raulet DH. Roles of natural killer cells in immunity to cancer, and applications to immunotherapy. Nat Rev Immunol. 2022;23(2):90–105.

12. Sivori S, Vacca P, Del Zotto G, Munari E, Mingari MC, Moretta L. Human NK cells: surface receptors, inhibitory checkpoints, and translational applications. Cell Mol Immunol. 2019;16(5):430.

13. Prager I, Watzl C. Mechanisms of natural killer cell-mediated cellular cytotoxicity. J Leukoc Biol. 2019;105(6):1319–29.

14. Ramírez-Labrada A, Pesini C, Santiago L, Hidalgo S, Calvo-Pérez A, Oñate C, et al. All About (NK Cell-Mediated) Death in Two Acts and an Unexpected Encore: Initiation, Execution and Activation of Adaptive Immunity. Front Immunol. 2022;13:896228.

15. Ochoa MC, Minute L, Rodriguez I, Garasa S, Perez-Ruiz E, Inogés S, et al. Antibody-dependent cell cytotoxicity: immunotherapy strategies enhancing effector NK cells. Immunol Cell Biol. 2017;95(4):347–55.

16. Alexandrov LB, Nik-Zainal S, Wedge DC, Aparicio SAJR, Behjati S, Biankin A V., et al. Signatures of mutational processes in human cancer. Nature. 2013;500(7463):415–21.

17. Fares J, Davis ZB, Rechberger JS, Toll SA, Schwartz JD, Daniels DJ, et al. Advances in NK cell therapy for brain tumors. npj Precis Oncol. 2023;7(1):1–17.

18. Morimoto T, Nakazawa T, Maeoka R, Nakagawa I, Tsujimura T, Matsuda R. Natural Killer Cell-Based Immunotherapy against Glioblastoma. Int J Mol Sci. 2023;24(3):2111.

19. Breznik B, Ko MW, Tse C, Chen PC, Senjor E, Majc B, et al. Infiltrating natural killer cells bind, lyse and increase chemotherapy efficacy in glioblastoma stem-like tumorospheres. Commun Biol. 2022;5(1).

20. Poorva P, Mast J, Cao B, Shah M V., Pollok KE, Shen J. Killing the killers: Natural killer cell therapy targeting glioma stem cells in high-grade glioma. Mol Ther. 2025;33(6):2462–78.

21. da Silva LHR, Catharino LCC, da Silva VJ, Evangelista GCM, Barbuto JAM. The War Is on: The Immune System against Glioblastoma—How Can NK Cells Drive This Battle? Biomedicines. 2022;10(2):400.

22. Zhong QY, Fan EX, Feng GY, Chen QY, Gou XX, Yue GJ, et al. A gene expression-based study on immune cell subtypes and glioma prognosis. BMC Cancer. 2019;19(1).

23. Crane CA, Han SJ, Barry JJ, Ahn BJ, Lanier LL, Parsa AT. TGF-beta downregulates the activating receptor NKG2D on NK cells and CD8+ T cells in glioma patients. Neuro Oncol. 2010;12(1):7– 13.

24. Friese MA, Wischhusen J, Wick W, Weiler M, Eisele G, Steinle A, et al. RNA interference targeting transforming growth factor-beta enhances NKG2D-mediated antiglioma immune response, inhibits glioma cell migration and invasiveness, and abrogates tumorigenicity in vivo. Cancer Res. 2004;64(20):7596–603.

25. Shaim H, Shanley M, Basar R, Daher M, Gumin J, Zamler DB, et al. Targeting the αv integrin/TGF-β axis improves natural killer cell function against glioblastoma stem cells. J Clin Invest. 2021;131(14):e142116.

26. Zhang Y, Guo F, Wang Y. Hypoxic tumor microenvironment: Destroyer of natural killer cell function. Chinese J Cancer Res. 2024;36(2):138.

27. Schindelin J, Arganda-Carreras I, Frise E, Kaynig V, Longair M, Pietzsch T, et al. Fiji: an open-source platform for biological-image analysis. Nat Methods. 2012;9(7):676–82.

28. Arora S, Singh S, Mittal A, Desai N, Khatri DK, Gugulothu D, et al. Spheroids in cancer research: Recent advances and opportunities. J Drug Deliv Sci Technol. 2024;100:106033.

29. Garcés-Lázaro I, Kotzur R, Cerwenka A, Mandelboim O. NK Cells Under Hypoxia: The Two Faces of Vascularization in Tumor and Pregnancy. Front Immunol. 2022;13:924775.

30. Vito A, El-Sayes N, Mossman K. Hypoxia-Driven Immune Escape in the Tumor Microenvironment. Cells. 2020;9(4):992.

31. Arner EN, Rathmell JC. Metabolic Programming and Immune Suppression in the Tumor Microenvironment. Cancer Cell. 2023;41(3):421.

32. Lobel GP, Jiang Y, Simon MC. Tumor microenvironmental nutrients, cellular responses, and cancer. Cell Chem Biol. 2023;30(9):1015–32.

33. Boucherit N, Gorvel L, Olive D. 3D Tumor Models and Their Use for the Testing of Immunotherapies. Front Immunol. 2020;11:603640.

34. He W, He W, Kuang Y, Xing X, Simpson RJ, Huang H, et al. Proteomic comparison of 3D and 2D glioma models reveals increased HLA-E expression in 3D models is associated with resistance to NK cell-mediated cytotoxicity. J Proteome Res. 2014;13(5):2272–81.

35. Braun FK, Rothhammer-Hampl T, Lorenz J, Pohl S, Menevse AN, Vollmann-Zwerenz A, et al. Scaffold-Based (MatrigelTM) 3D Culture Technique of Glioblastoma Recovers a Patient-like Immunosuppressive Phenotype. Cells. 2023;12(14).

36. Burster T, Gärtner F, Bulach C, Zhanapiya A, Gihring A, Knippschild U. Regulation of MHC I Molecules in Glioblastoma Cells and the Sensitizing of NK Cells. Pharmaceuticals. 2021;14(3):236.

37. Robilliard LD, Yu J, Anchan A, Finlay G, Angel CE, Graham ES. Comprehensive Assessment of Secreted Immuno-Modulatory Cytokines by Serum-Differentiated and Stem-like Glioblastoma Cells Reveals Distinct Differences between Glioblastoma Phenotypes. Int J Mol Sci. 2022;23(22):14164.

38. Chang AL, Miska J, Wainwright DA, Dey M, Rivetta C V., Yu D, et al. CCL2 produced by the glioma microenvironment is essential for the recruitment of regulatory t cells and myeloid-derived suppressor cells. Cancer Res. 2016;76(19):5671–82.

39. Takacs GP, Kreiger CJ, Luo D, Tian G, Garcia JS, Deleyrolle LP, et al. Glioma-derived CCL2 and CCL7 mediate migration of immune suppressive CCR2+/CX3CR1+ M-MDSCs into the tumor microenvironment in a redundant manner. Front Immunol. 2023;13:993444.

40. Vakilian A, Khorramdelazad H, Heidari P, Sheikh Rezaei Z, Hassanshahi G. CCL2/CCR2 signaling pathway in glioblastoma multiforme. Neurochem Int. 2017;103:1–7.

41. Alfaro C, Teijeira A, Oñate C, Perez G, Sanmamed MF, Andueza MP, et al. Tumor-Produced Interleukin-8 Attracts Human Myeloid-Derived Suppressor Cells and Elicits Extrusion of Neutrophil Extracellular Traps (NETs). Clin Cancer Res. 2016;22(15):3924–36.

42. Jackson C, Cherry C, Bom S, Dykema AG, Wang R, Thompson E, et al. Distinct myeloid-derived suppressor cell populations in human glioblastoma. Science. 2025;387(6731).

43. Weber R, Groth C, Lasser S, Arkhypov I, Petrova V, Altevogt P, et al. IL-6 as a major regulator of MDSC activity and possible target for cancer immunotherapy. Cell Immunol. 2021;359.

44. Morrison BE, Park SJ, Mooney JM, Mehrad B. Chemokine-mediated recruitment of NK cells is a critical host defense mechanism in invasive aspergillosis. J Clin Invest. 2003;112(12):1862.

45. He Q, Shi X, Zhou B, Teng J, Zhang C, Liu S, et al. Interleukin 8 (CXCL8)-CXC chemokine receptor 2 (CXCR2) axis contributes to MiR-4437-associated recruitment of granulocytes and natural killer cells in ischemic stroke. Mol Immunol. 2018;101:440–9.

46. Vujanovic L, Ballard W, Thorne SH, Vujanovic NL, Butterfield LH. Adenovirus-engineered human dendritic cells induce natural killer cell chemotaxis via CXCL8/IL-8 and CXCL10/IP-10. Oncoimmunology. 2012;1(4):448–57.

47. Serrano-Pertierra E, Blanco-Gelaz MA, Oliva-Nacarino P, Martínez-Camblor P, Villafani J, López-Larrea C, et al. Increased natural killer cell chemotaxis to CXCL12 in patients with multiple sclerosis. J Neuroimmunol. 2015;282:39–44.

48. Ran G he, Lin Y qing, Tian L, Zhang T, Yan D mei, Yu J hua, et al. Natural killer cell homing and trafficking in tissues and tumors: from biology to application. Signal Transduct Target Ther. 2022;7(1):1–21.

49. Chan TYH, Wong JSY, Kiang KMY, Sun CWY, Leung GKK. The duality of CXCR3 in glioblastoma: unveiling autocrine and paracrine mechanisms for novel therapeutic approaches. Cell Death Dis. 2023;14(12):1–13.

50. Avril T, Vauleon E, Hamlat A, Saikali S, Etcheverry A, Delmas C, et al. Human glioblastoma stem-like cells are more sensitive to allogeneic NK and T cell-mediated killing compared with serum-cultured glioblastoma cells. Brain Pathol. 2012;22(2):159–74.

51. Haspels HN, Rahman MA, Joseph JV, Navarro AG, Chekenya M. Glioblastoma stem-like cells are more susceptible than differentiated cells to natural killer cell lysis mediated through killer immunoglobulin-like receptors-human leukocyte antigen ligand mismatch and activation receptor-ligand interactions. Front Immunol. 2018;9:1345.

52. Close HJ, Stead LF, Nsengimana J, Reilly KA, Droop A, Wurdak H, et al. Expression profiling of single cells and patient cohorts identifies multiple immunosuppressive pathways and an altered NK cell phenotype in glioblastoma. Clin Exp Immunol. 2020;200(1):33.

53. Karimi E, Yu MW, Maritan SM, Perus LJM, Rezanejad M, Sorin M, et al. Single-cell spatial immune landscapes of primary and metastatic brain tumours. Nat. 2023;614(7948):555–63.

54. Fu W, Wang W, Li H, Jiao Y, Huo R, Yan Z, et al. Single-Cell Atlas Reveals Complexity of the Immunosuppressive Microenvironment of Initial and Recurrent Glioblastoma. Front Immunol. 2020;11:835.

55. Wu J, Gao FX, Wang C, Qin M, Han F, Xu T, et al. IL-6 and IL-8 secreted by tumour cells impair the function of NK cells via the STAT3 pathway in oesophageal squamous cell carcinoma. J Exp Clin Cancer Res. 2019;38(1):1–15.

56. Han B, Mao FY, Zhao YL, Lv YP, Teng YS, Duan M, et al. Altered NKp30, NKp46, NKG2D, and DNAM-1 Expression on Circulating NK Cells Is Associated with Tumor Progression in Human Gastric Cancer. J Immunol Res. 2018;2018:6248590.

57. Mamessier E, Pradel LC, Thibult M-L, Drevet C, Zouine A, Jacquemier J, et al. Peripheral blood NK cells from breast cancer patients are tumor-induced composite subsets. J Immunol. 2013;190(5):2424–36.

58. Molfetta R, Zitti B, Lecce M, Milito ND, Stabile H, Fionda C, et al. CD155: A Multi-Functional Molecule in Tumor Progression. Int J Mol Sci. 2020;21(3):922.

59. Zhan M, Zhang Z, Zhao X, Zhang Y, Liu T, Lu L, et al. CD155 in tumor progression and targeted therapy. Cancer Lett. 2022;545:215830.

60. Sloan KE, Eustace BK, Stewart JK, Zehetmeier C, Torella C, Simeone M, et al. CD155/PVR plays a key role in cell motility during tumor cell invasion and migration. BMC Cancer. 2004;4:73.

61. Sloan KE, Stewart JK, Treloar AF, Matthews RT, Jay DG. CD155/PVR enhances glioma cell dispersal by regulating adhesion signaling and focal adhesion dynamics. Cancer Res. 2005;65(23):10930–7.

62. Enloe BM, Jay DG. Inhibition of Necl-5 (CD155/PVR) reduces glioblastoma dispersal and decreases MMP-2 expression and activity. J Neurooncol. 2011;102(2):225–35.

63. Yeo J, Ko M, Lee DH, Park Y, Jin HS. TIGIT/CD226 Axis Regulates Anti-Tumor Immunity. Pharmaceuticals (Basel). 2021;14(3):1–20.

64. Lupo KB, Torregrosa-Allen S, Elzey BD, Utturkar S, Lanman NA, Cohen-Gadol AA, et al. TIGIT contributes to the regulation of 4-1BB and does not define NK cell dysfunction in glioblastoma. iScience. 2023;26(12):108353.

