## Supplementary File for "Glioblastoma–natural killer cell crosstalk: insights from dynamic spheroid models reveal the importance of secreted cytokines and the CD155 axis"

Table S1: Antibodies used for immunofluorescent staining of FFPE slides.

| Group | Antibody | Catalog number | Company | Dilution |
|---|---|---|---|---|
| Primary antibodies | Mouse anti-human CD45 antibody, clone 2D1 | MAB1430 | R&D Systems | 1:50 |
| | Rabbit anti-GFAP antibody, polyclonal | ab211271 | abcam | 1:1,000 |
| | Mouse anti-human SOX2, clone 20G5 | ab171380 | abcam | 1:50 |
| | Recombinant rabbit anti-OLIG2 antibody, clone EPR2673 | ab109186 | abcam | 1:100 |
| | Rabbit anti-Ki67 antibody, polyclonal | ab15580 | abcam | 1:1,000 |
| | Mouse anti-human HIF1A antibody, clone mgc3 | MA1-516 | Thermo Fisher Scientific | 1:100 |
| | Mouse anti-human CD44 antibody, clone Bu52 | MCA2504 | Bio-Rad | 1:100 |
| | Rabbit anti-Carbonic Anhydrase 9 (CA9), clone EPR23055-5 | ab243660 | abcam | 1:200 |
| Secondary antibodies | Goat anti-rabbit IgG (H+L) highly cross-adsorbed secondary antibody, Alexa fluor plus 488 | A11008 | Thermo Fisher Scientific | 1:200 |
| | Goat anti-mouse IgG (H+L) highly cross-adsorbed secondary antibody, Alexa fluor plus 546 | A11003 | Thermo Fisher Scientific | 1:200 |

Table S2: Antibodies used for flow cytometry.

| Group | Antibody | Catalog number | Company | Dilution |
|---|---|---|---|---|
| Receptors on NK cells | APC anti-human CD158e1 (KIR3DL1, NKB1) Antibody, clone DX9 | 312715 | BioLegend | 1:20 |
| | APC anti-human CD226 (DNAM-1) Antibody, clone 11A8 | 338311 | BioLegend | 1:20 |
| | APC anti-human CD337 (NKp30) Antibody, clone P30-15 | 325209 | BioLegend | 1:20 |
| | APC anti-human CD96 (TACTILE) Antibody, clone NK-92.39 | 338409 | BioLegend | 1:20 |
| | FITC anti-human CD158b/j (KIR2DL2/L3/S2) Antibody, clone DX27 | 312603 | BioLegend | 1:20 |
| | FITC anti-human CD16 Antibody, clone 3G8 | 302005 | BioLegend | 1:20 |
| | FITC anti-human CD279 (PD-1) Antibody, clone EH12.2H7 | 329903 | BioLegend | 1:20 |
| | FITC anti-human CD94 Antibody, clone DX22 | 305504 | BioLegend | 1:20 |
| | PE anti-human CD158a (KIR2DL1) Antibody, clone HP-DM1 | 374903 | BioLegend | 1:20 |
| | PE anti-human CD158d (KIR2DL4) Antibody, clone mAb 33 (33) | 347005 | BioLegend | 1:20 |
| | PE anti-human CD314 (NKG2D) Antibody. Clone 1D11 | 320805 | BioLegend | 1:20 |
| | PE anti-human TIGIT (VSTM3) Antibody, clone A15153G | 372703 | BioLegend | 1:20 |
| Ligands for NK cell receptors | Alexa Fluor® 488 anti-human ULBP-1 Antibody, clone 170818 | FAB1380G | R&D Systems | 1:20 |
| | anti-human ULBP-3 Antibody, clone 166510 | MAB1517-SP | R&D Systems | 1:20 |
| | APC anti-human CD112 (Nectin-2) Antibody, clone TX31 | 337411 | BioLegend | 1:20 |
| | APC anti-human HLA-E Antibody, clone 3D12 | 342605 | BioLegend | 1:20 |
| | APC anti-human MICA/MICB Antibody, clone 6D4 | 320907 | BioLegend | 1:20 |
| | APC anti-human ULBP-2/5/6 Antibody, clone 165903 | FAB1298A | R&D Systems | 1:10 |
| | FITC anti-human CD155 (PVR) Antibody, clone SKII.4 | 337627 | BioLegend | 1:20 |
| | PE anti human B7-H6 Antibody, clone JAM1EW | 12-6526-42 | Thermo Fisher Scientific | 1:20 |
| | PE anti-human CD274 (B7-H1, PD-L1) Antibody, clone 29E.2A3 | 329705 | BioLegend | 1:20 |
| | PE anti-human CD54 Antibody, clone HA58 | 353105 | BioLegend | 1:20 |

| | | | | |
|---|---|---|---|---|
| | PE anti-human HLA-A,B,C Antibody, clone W6/32 | 311405 | BioLegend | 1:20 |
| Isotype controls | APC Mouse IgG1, κ Isotype Ctrl (FC) Antibody, clone MOPC-21 | 400121 | BioLegend | Assay dependent |
| | APC Mouse IgG2a, κ Isotype Ctrl Antibody, clone MOPC-173 | 400219 | BioLegend | Assay dependent |
| | FITC Mouse IgG1, κ Isotype Ctrl (FC) Antibody, clone MOPC-21 | 400109 | BioLegend | Assay dependent |
| | FITC Mouse IgG2a, κ Isotype Ctrl (FC) Antibody, clone MOPC-173 | 400209 | BioLegend | Assay dependent |
| | PE Mouse IgG1, κ Isotype Ctrl (FC) Antibody, clone MOPC-21 | 400113 | BioLegend | Assay dependent |
| | PE Mouse IgG2a, κ Isotype Ctrl (FC) Antibody, clone MOPC-173 | 400213 | BioLegend | Assay dependent |
| | PE Mouse IgG2b, κ Isotype Ctrl Antibody, clone MPC-11, clone MPC-11 | 400313 | BioLegend | Assay dependent |
| | Purified Mouse IgG2a, κ Isotype Ctrl Antibody, clone MG2a-53 | 401501 | BioLegend | Assay dependent |

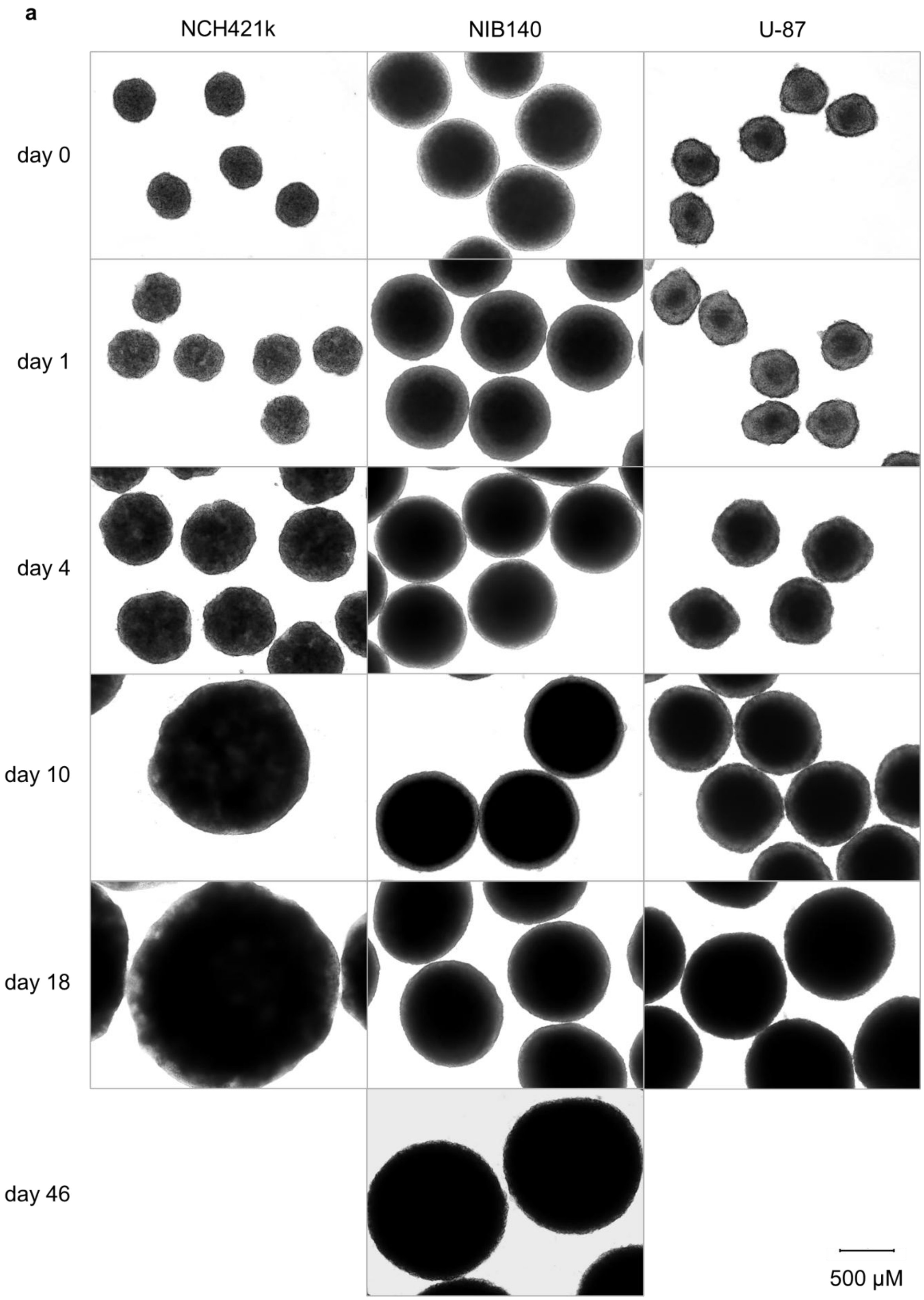


Figure S1: **Growth of GB spheroids in the Celvivo system.** Representative images of NCH421k, NIB140 and U-87 spheroids at selected time points are shown.

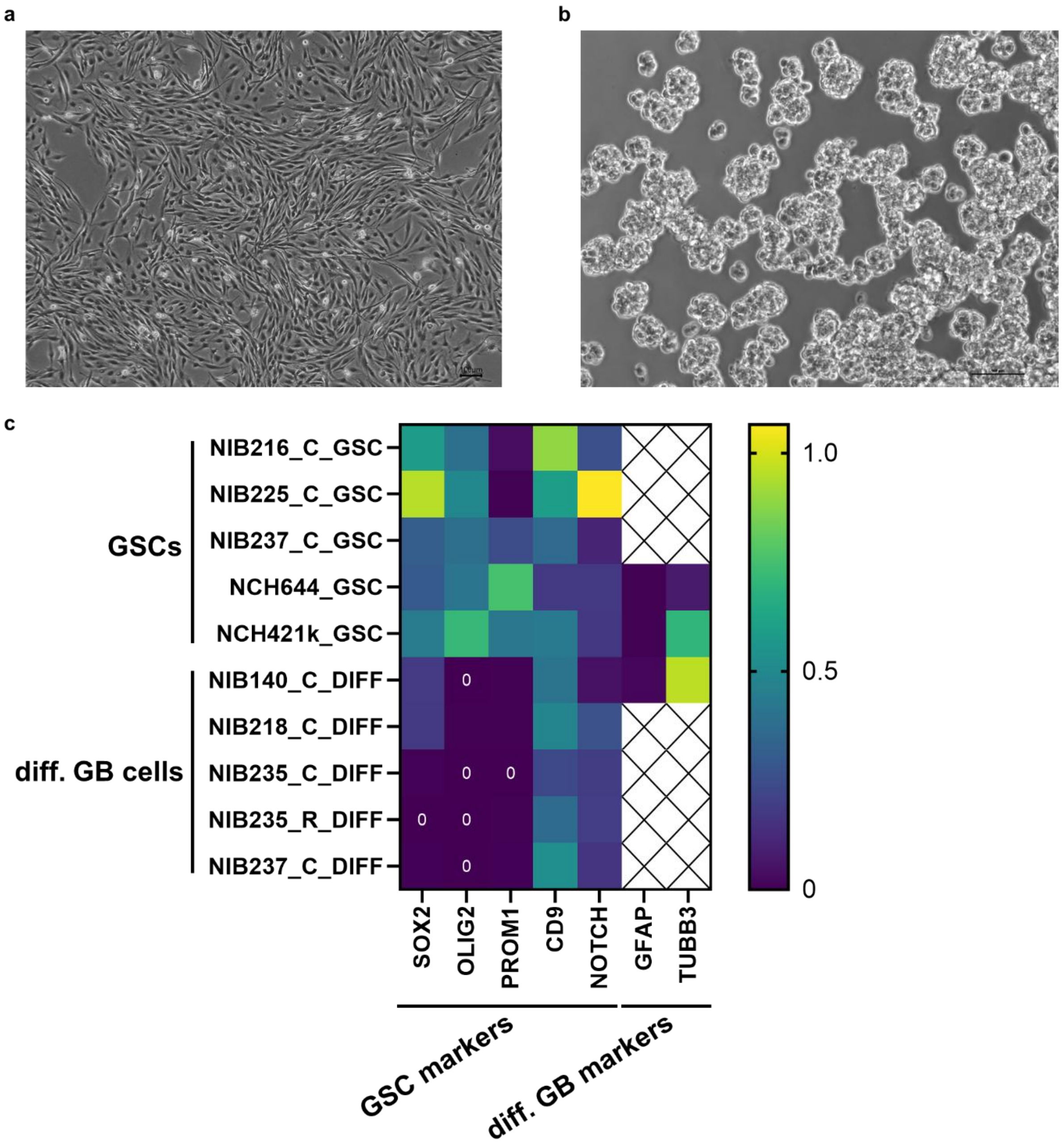


Figure S2: **Comparison of cell lines NCH421k and NIB140.** a) NCH421k cells are standardly cultured in suspension as small multicellular spheres. b) NIB140 is an adherent cell line. c) Comparison of the expression of stemness-related marker genes (SOX2, OLIG2, PROM1, CD9, NOTCH) and markers of differentiated GB cells (GFAP, TUBB3) in several GSC cell lines and cell lines of differentiated GB cells. mRNA expression of selected marker genes were determined by RT-qPCR, normalized to the expression of housekeeping genes HPRT1 and GAPDH, and analyzed with quantGenius software. Data were re-analysed from Majc et al., 2022 and Majc et al., 2024.

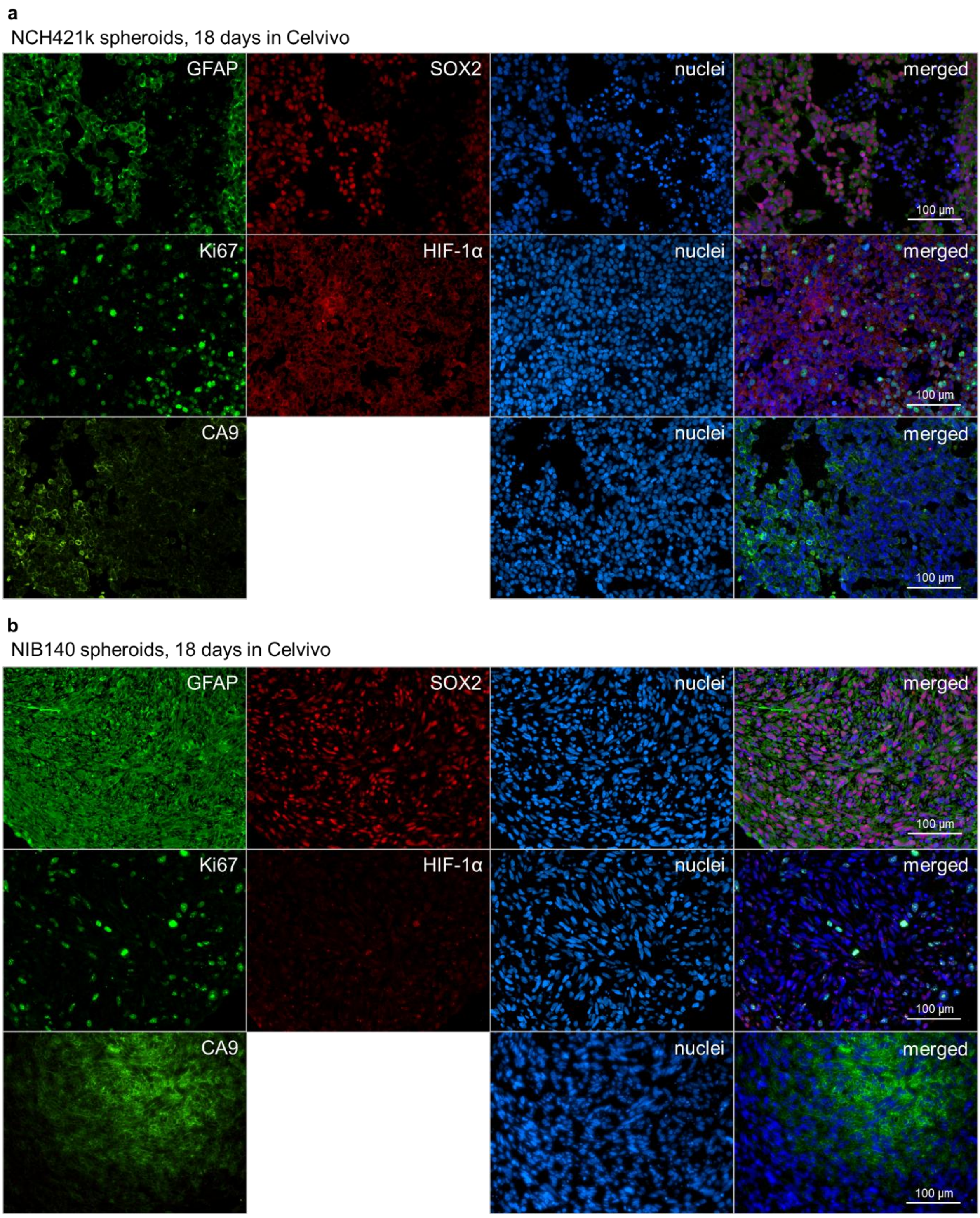


Figure S3: **Characterisation of GB spheroids cultured in the Celvivo system for 18 days.** Immunofluorescent staining of differentiation marker GFAP, stemness marker SOX2, proliferation marker Ki-67 and hypoxia markers HIF-1α and CA9 in NCH421k spheroids (a) and NIB140 spheroids (b).

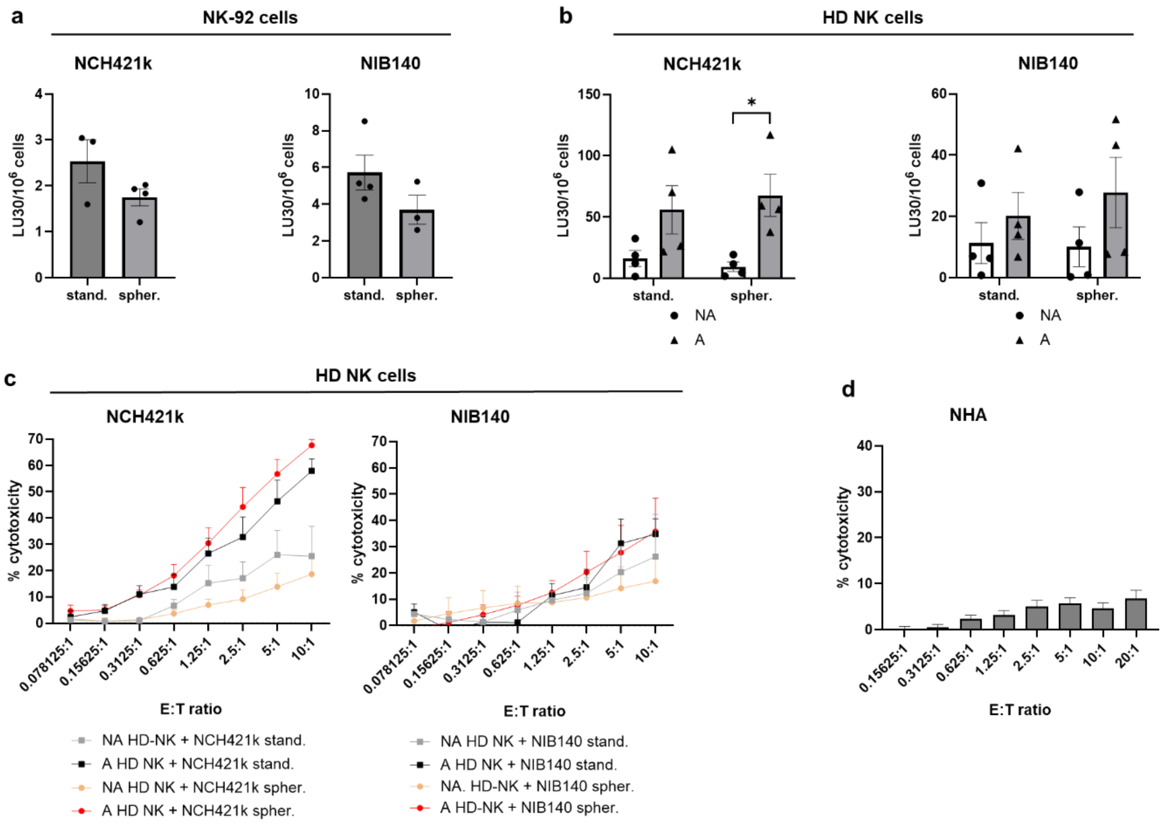


Figure S4: **Cytotoxicity of NK cells against single cells dissociated from NCH421k and NIB140 spheroids**. LU30 as determined by the calcein release assay of NK-92 cells (a) and HD NK cells (b) against dissociated NCH421k and NIB140 spheroids and NCH421k and NIB140 cells from the standard culture. Data are presented as mean ± SEM of 3-4 biological replicates, depicted as individual spots. Unpaired t-test (a) and 2-way ANOVA with uncorrected Fisher's least significant difference (b) were used for statistical analysis. c) % cytotoxicity plots of calcein release assay of HD NK cells against dissociated NCH421k and NIB140 spheroids and NCH421k and NIB140 cells from the standard culture. Data are presented as mean ± SEM of 3-4 biological replicates. 2-way ANOVA with Tukey's multiple comparisons test was used for statistical analysis. NA – nonactivated cells, A – IL2-activated cells. d) NK-92 cell cytotoxicity against NHA as measured by the calcein release assay. Data are presented as mean ± SEM of 4 biological replicates.

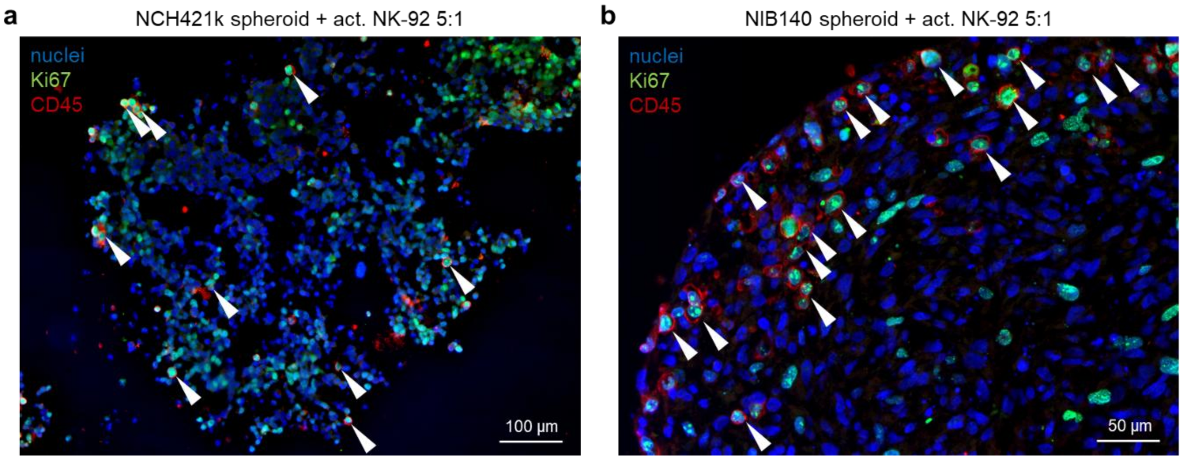


Figure S5: **NK-92 cells proliferate in the GB spheroids.** FFPE sections of NCH421k (a) and NIB140 (b) cocultures with NK-92 cells were stained to detect CD45-positive NK-92 cells and Ki67-positive proliferating cells. Proliferating NK-92 cells are indicated with white arrows.

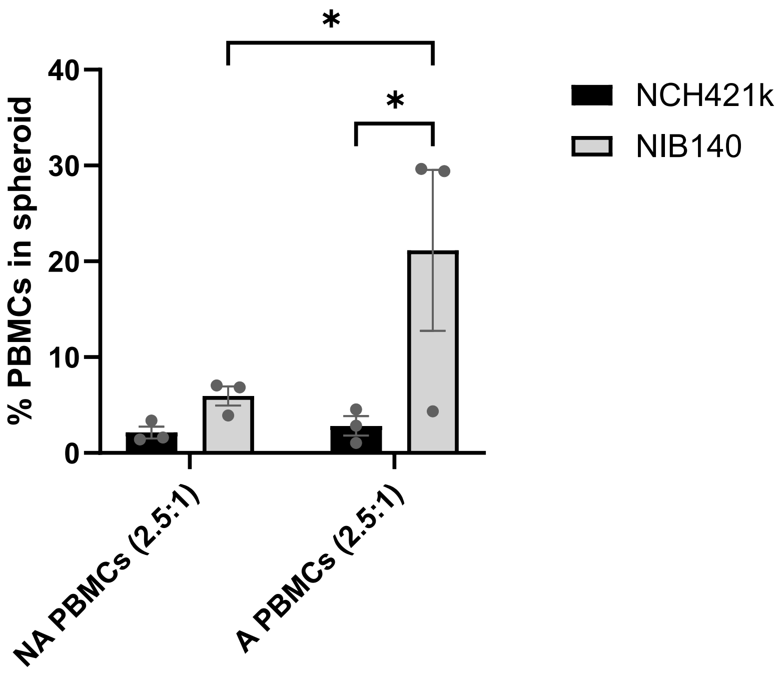


Figure S6: **PBMCs from GB patients infiltrate at higher levels into NIB140 spheroids compared to NCH421k spheroids.** After 24h of direct co-culture of GB spheroids and PBMCs, infiltrated pre-labelled PBMCs were detected by flow cytometry. The experiment was performed with PBMCs from three GB patient PBMCs. Data are presented as mean ± SEM and data points represent biological replicates.

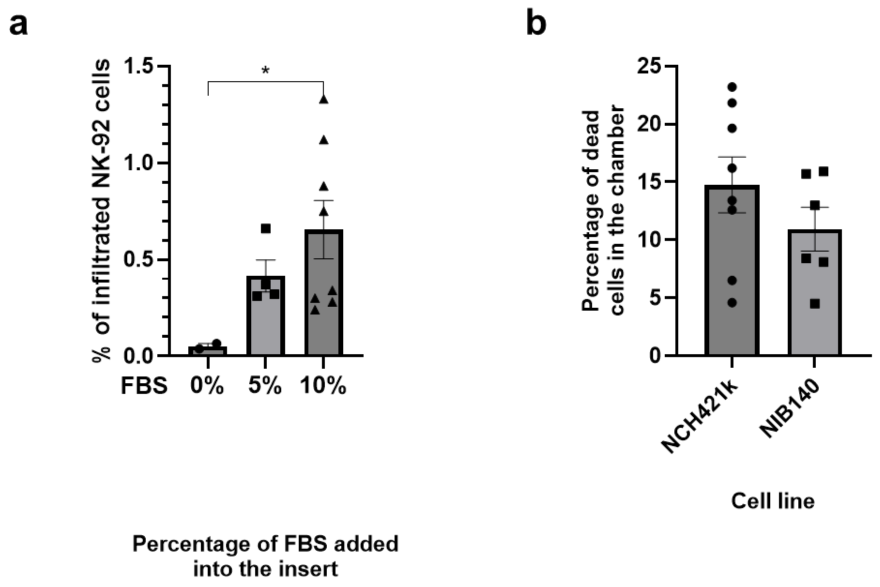


Figure S7: **Additional results on NK-92 cell infiltration and cytotoxicity against GB spheroids tested on the organ-on-chip platform**. **a)** The effect of FBS in the chamber on the infiltration of NK-92 cells. Data are presented as mean ± SEM and data points represent each of the 2-8 biological replicates. Brown-Forsythe and Welch ANOVA was used for statistical analysis. **b)** Percentage of dead GB cells in the chamber. Data are presented as mean ± SEM and data points represent each of the 6-8 biological replicates. Unpaired t-test was used for statistical analysis.

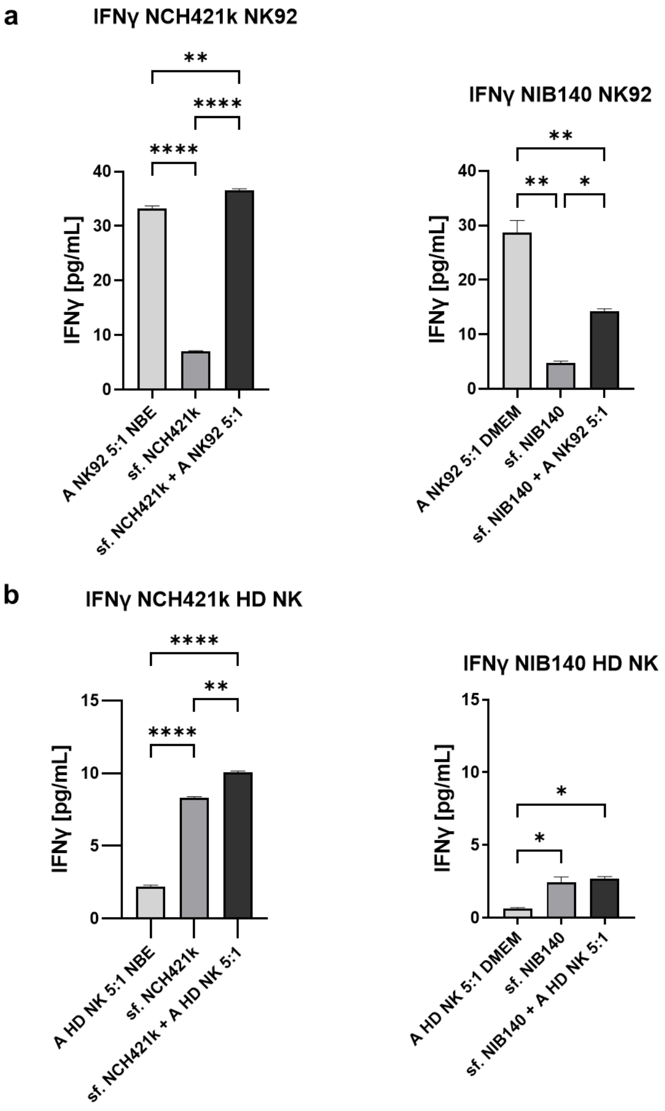


Figure S8: **IFN-γ levels in the media of NK-92 (a) and HD NK cell (b) cocultures with NCH421k and NIB140 spheroids**. Data are presented as mean ± SEM of 2 technical replicates. Ordinary one-way ANOVA was used for statistical analysis.

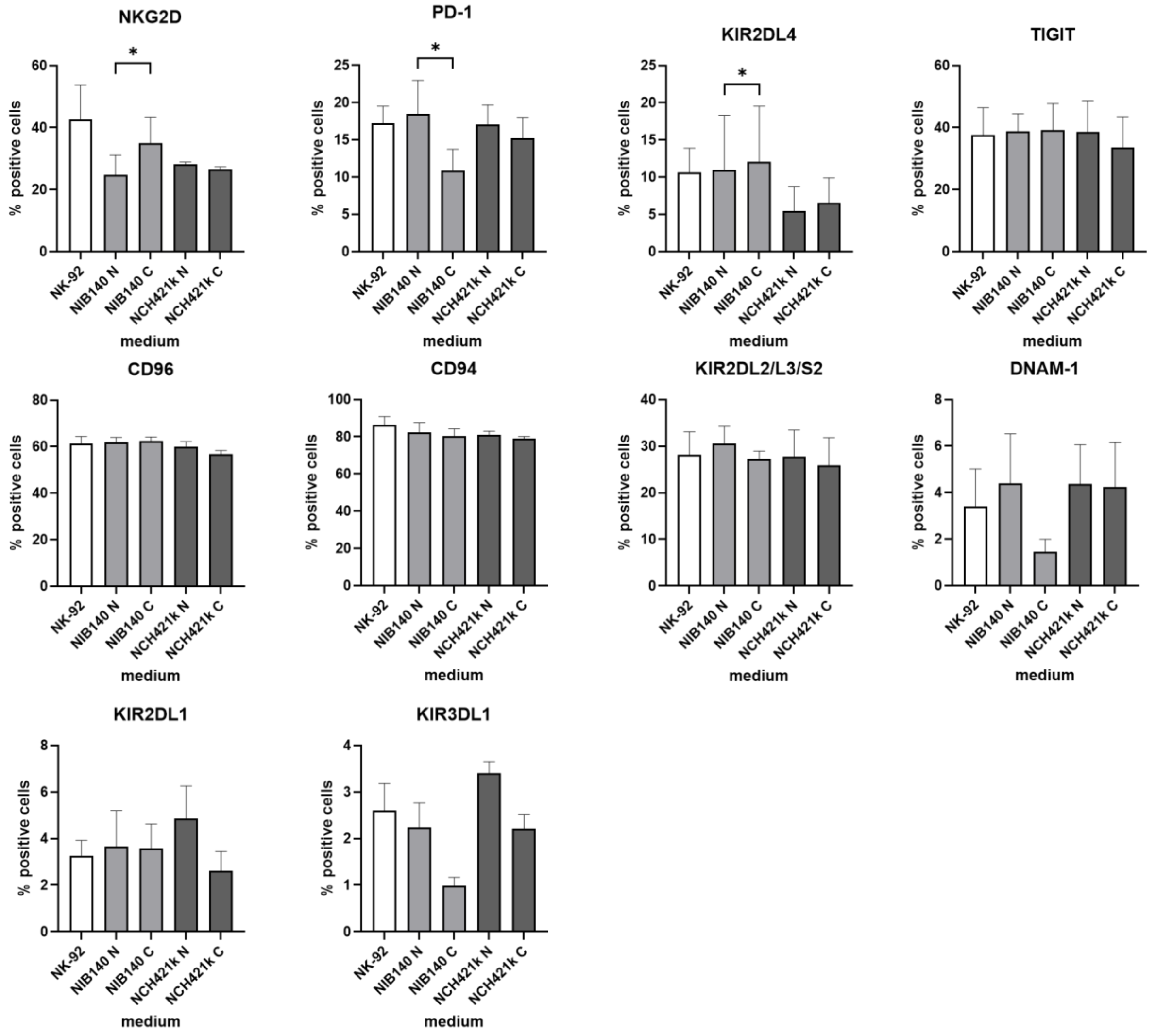


Figure S9: **Expression of NK cell receptors on NK-92 cells cultured in NCH421k- or NIB140-conditioned medium**. Data are presented as mean ± SEM of 3-4 biological replicates. Paired t-test was used for statistical analysis. N – nonconditioned medium, C – conditioned medium.

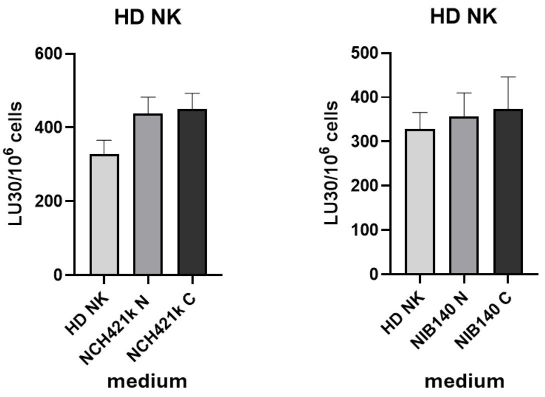


Figure S10: **The effect of NCH421k- and NIB140-conditioned medium on cytotoxicity of HD NK cells against K562 cells**. Data are presented as means ± SEM of 3 biological replicates. Paired t-test was used for statistical analysis. N – nonconditioned medium, C – conditioned medium.

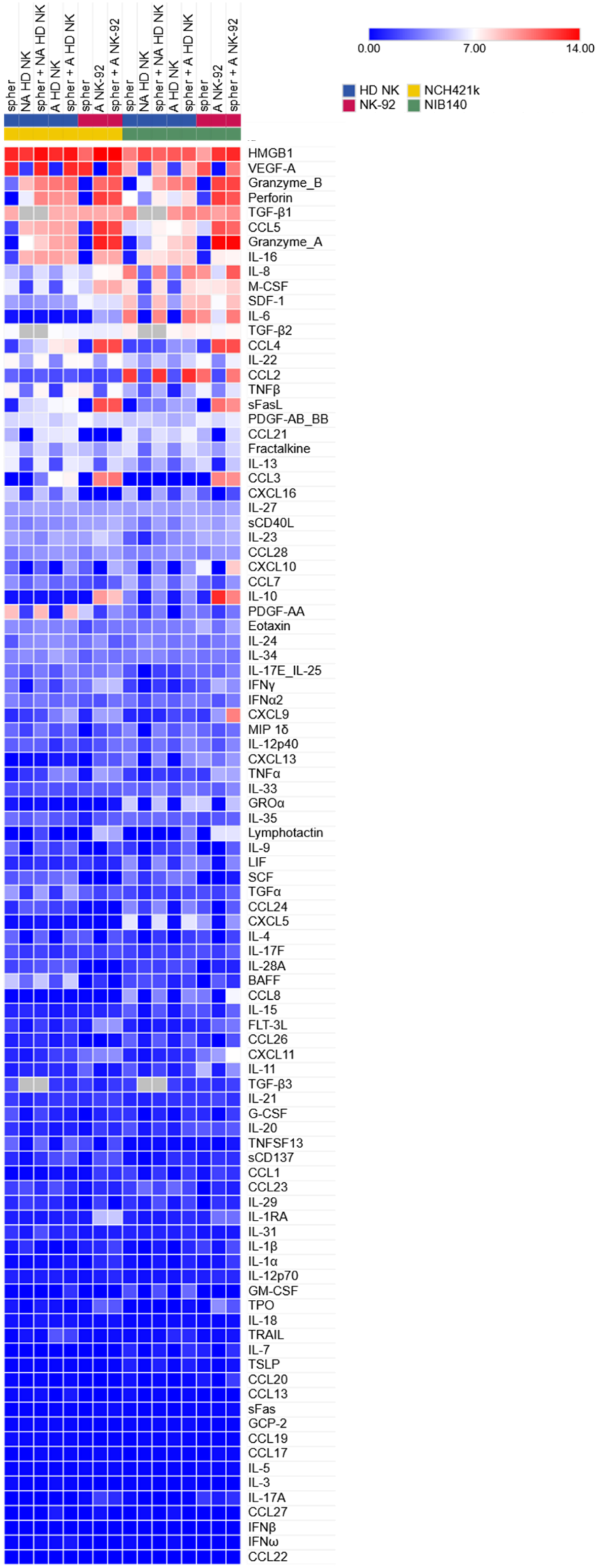
spher
NA HD NK
spher + NA HD NK
A HD NK
spher + A HD NK
spher
A NK-92
spher + A NK-92
spher
NA HD NK
spher + NA HD NK
A HD NK
spher + A HD NK
spher
A NK-92
spher + A NK-92
0.00
7.00
14.00
HD NK
NCH421k
NK-92
NIB140
HMGB1
VEGF-A
Granzyme_B
Perforin
TGF-β1
CCL5
Granzyme_A
IL-16
IL-8
M-CSF
SDF-1
IL-6
TGF-β2
CCL4
IL-22
CCL2
TNFβ
sFasL
PDGF-AB_BB
CCL21
Fractalkine
IL-13
CCL3
CXCL16
IL-27
sCD40L
IL-23
CCL28
CXCL10
CCL7
IL-10
PDGF-AA
Eotaxin
IL-24
IL-34
IL-17E_IL-25
IFNγ
IFNα2
CXCL9
MIP 1δ
IL-12p40
CXCL13
TNFα
IL-33
GROα
IL-35
Lymphotactin
IL-9
LIF
SCF
TGFα
CCL24
CXCL5
IL-4
IL-17F
IL-28A
BAFF
CCL8
IL-15
FLT-3L
CCL26
CXCL11
IL-11
TGF-β3
IL-21
G-CSF
IL-20
TNFSF13
sCD137
CCL1
CCL23
IL-29
IL-1RA
IL-31
IL-1β
IL-1α
IL-12p70
GM-CSF
TPO
IL-18
TRAIL
IL-7
TSLP
CCL20
CCL13
sFas
GCP-2
CCL19
CCL17
IL-5
IL-3
IL-17A
CCL27
IFNβ
IFNω
CCL22

Figure S11: **Analysis of secreted cytokines in the 24h spheroid-HD NK cell co-cultures**. log2(x+1)-transformed cytokine concentrations (averaged from two technical replicates) are presented in the heatmap.

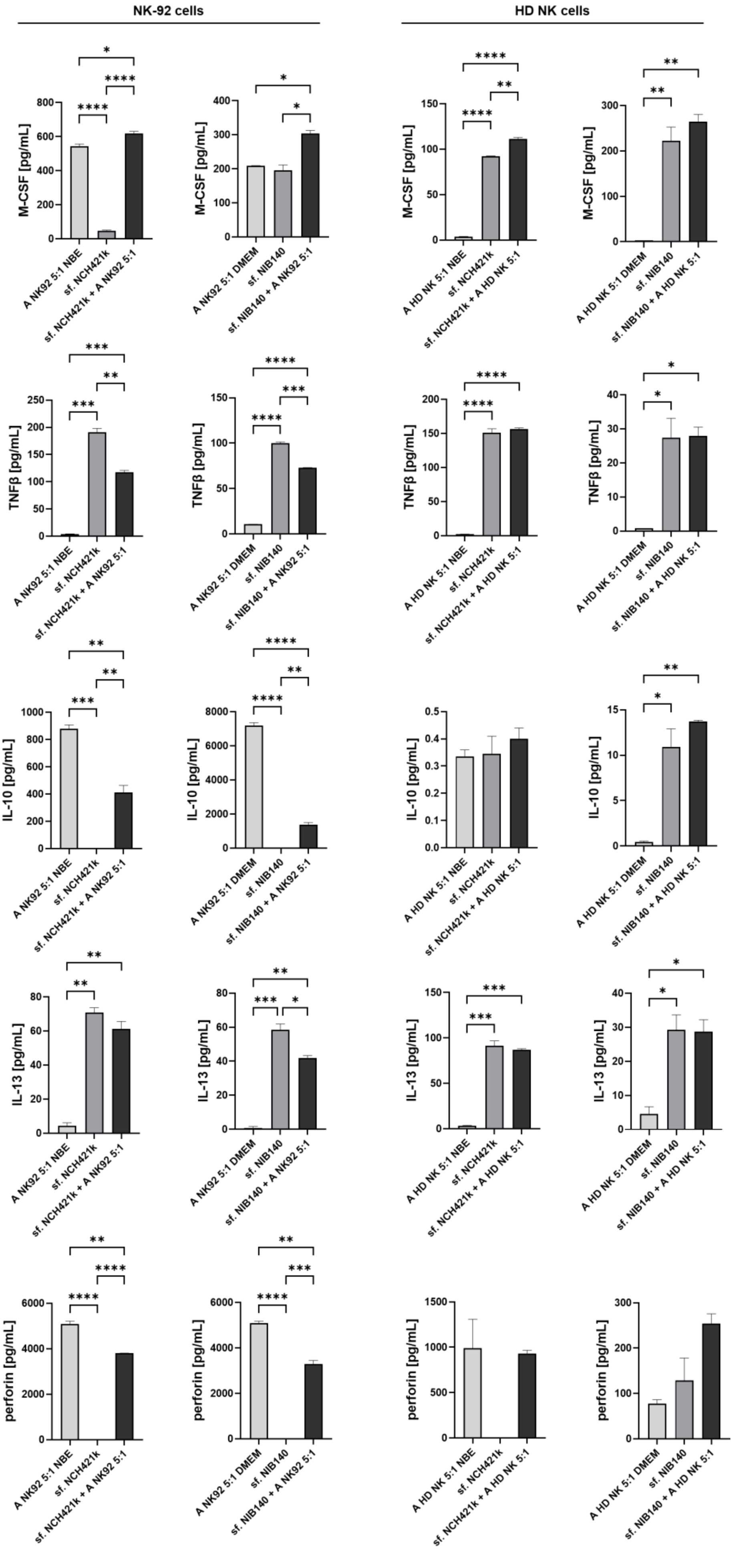
NK-92 cells
HD NK cells
M-CSF [pg/mL]
TNFβ [pg/mL]
IL-10 [pg/mL]
IL-13 [pg/mL]
perforin [pg/mL]
A NK92 5:1 NBE
sf. NCH421k
sf. NCH421k + A NK92 5:1
A NK92 5:1 DMEM
sf. NIB140
sf. NIB140 + A NK92 5:1
A HD NK 5:1 NBE
sf. NCH421k + A HD NK 5:1
A HD NK 5:1 DMEM
sf. NIB140 + A HD NK 5:1

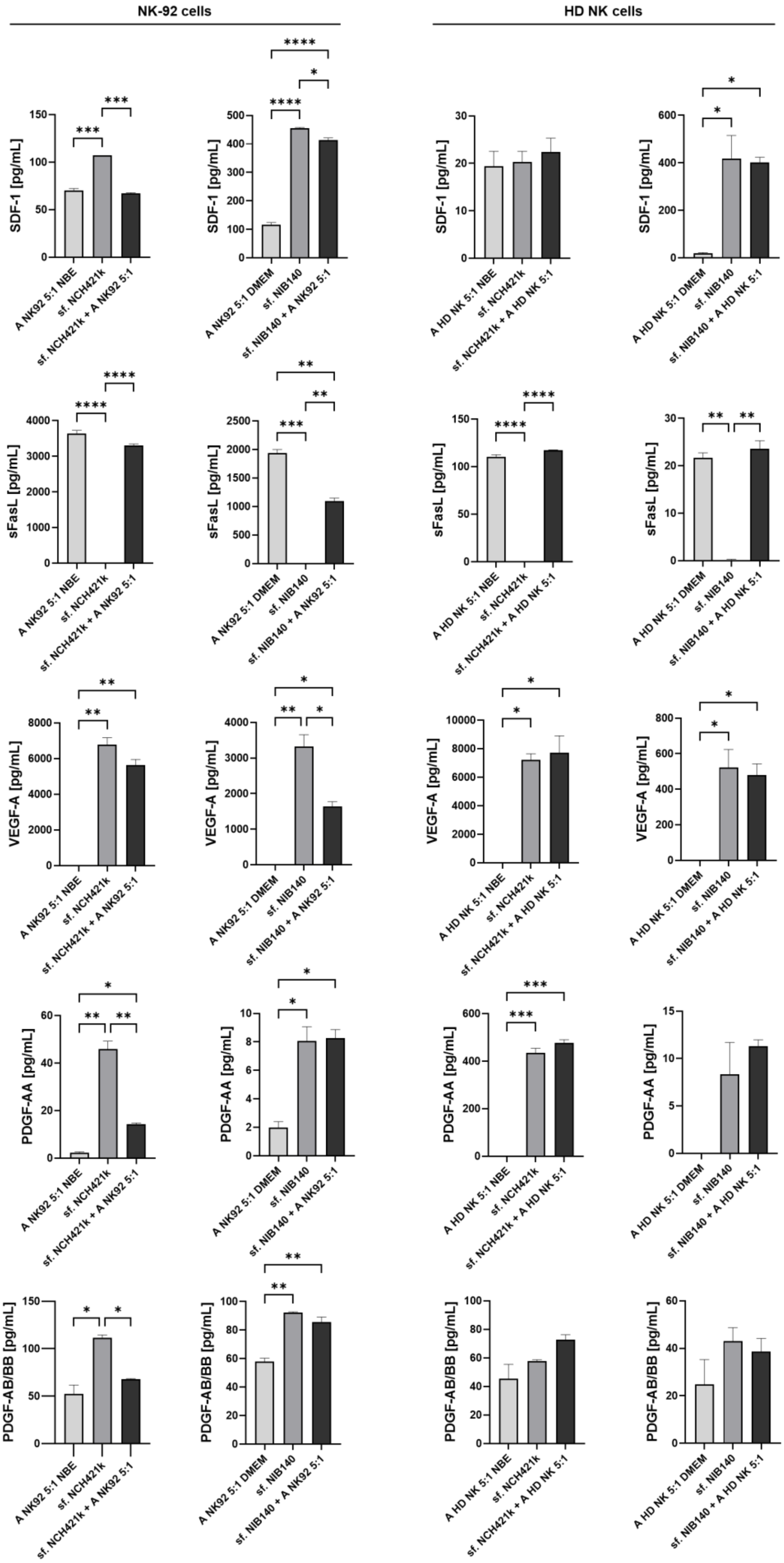
NK-92 cells
HD NK cells
SDF-1 [pg/mL]
sFasL [pg/mL]
VEGF-A [pg/mL]
PDGF-AA [pg/mL]
PDGF-AB/BB [pg/mL]
A NK92 5:1 NBE
sf. NCH421k
sf. NCH421k + A NK92 5:1
A NK92 5:1 DMEM
sf. NIB140
sf. NIB140 + A NK92 5:1
A HD NK 5:1 NBE
sf. NCH421k + A HD NK 5:1
A HD NK 5:1 DMEM
sf. NIB140 + A HD NK 5:1

Figure S12: **Secreted factors that were modulated in GB spheroid-NK cell co-cultures.** Data are presented as mean ± SEM of 2 technical replicates. Ordinary one-way ANOVA was used for statistical analysis.

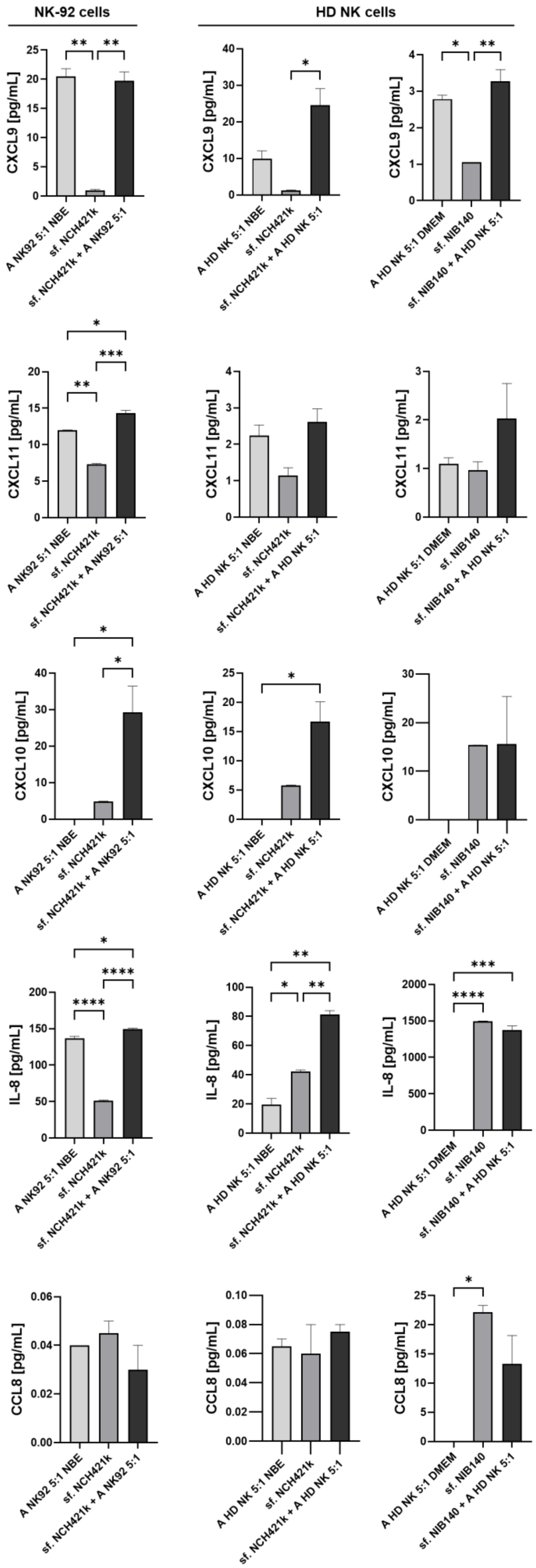
NK-92 cells
HD NK cells
CXCL9 [pg/mL]
CXCL11 [pg/mL]
CXCL10 [pg/mL]
IL-8 [pg/mL]
CCL8 [pg/mL]
A NK92 5:1 NBE
sf. NCH421k
sf. NCH421k + A NK92 5:1
A HD NK 5:1 NBE
sf. NCH421k + A HD NK 5:1
A HD NK 5:1 DMEM
sf. NIB140
sf. NIB140 + A HD NK 5:1

Figure S13: **Concentrations of secreted factors that were strikingly upregulated in NIB140 spheroid-NK-92 cell co-cultures in other GB-NK cell co-cultures**. Data are presented as mean ± SEM of 2 technical replicates. Ordinary one-way ANOVA was used for statistical analysis.

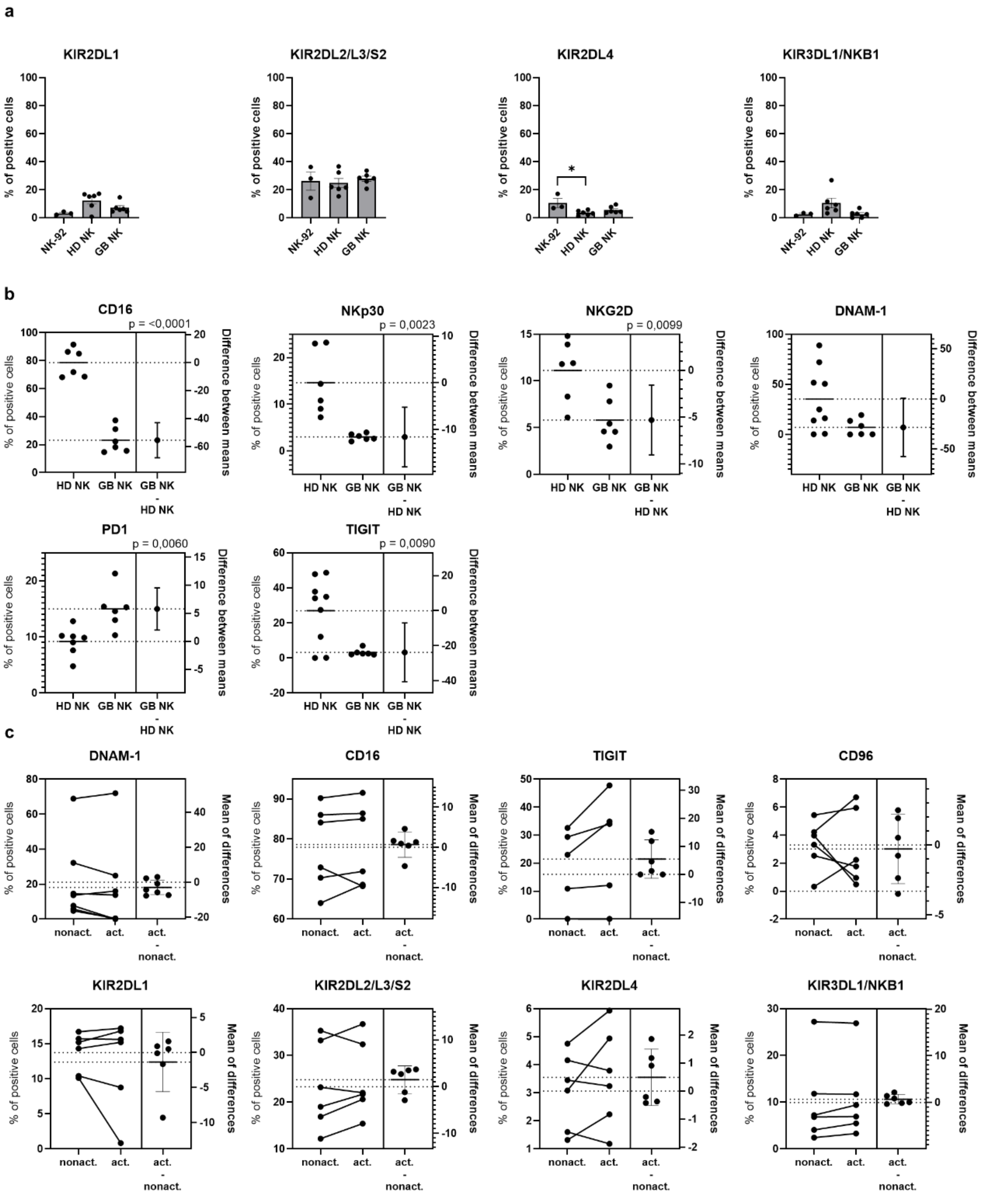


Figure S14: **Expression of NK cell receptors on IL2-activated NK-92 cells, HD NK cells and NK cells from GB patients.** a) Comparison of receptor expression on IL-2-activated NK-92 cells, HD NK cells (N=6) and NK cells from GB patients (N=6). Data are presented as mean ± SEM and dots represent biological replicates. Ordinary one-way ANOVA was used for statistical analysis. b) Estimation plots showing the expression of NK cell receptors, differentially expressed between HD (N=6-9) and GB NK cells (N=6). c) Estimation plots showing the expression of NK cell receptors on HD NK cells with and without IL-2 activation (N=6).

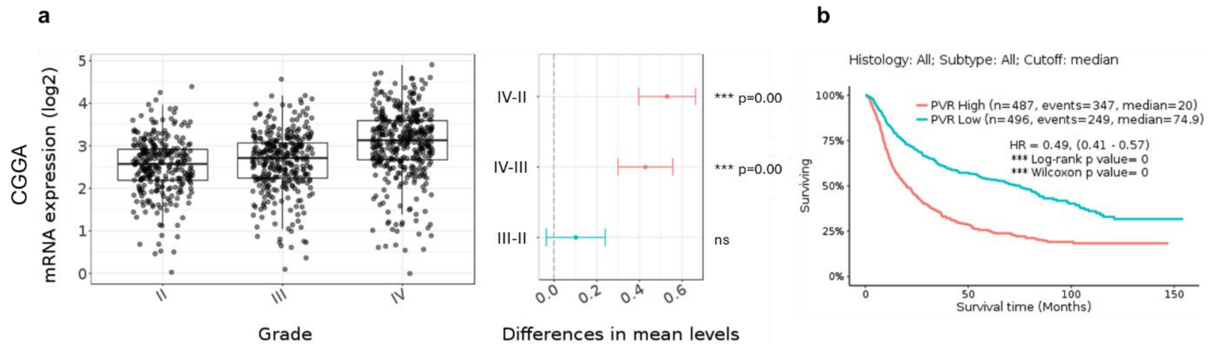


Figure S15: **Expression of *PVR* in glioma according to the CGGA data**. a) According to the TCGA data, *PVR* is upregulated in GB compared to lower-grade gliomas. Tukey's Honest Significant Difference were calculated in GlioVis. b) Higher expression of *PVR* is associated with shorter survival of glioma patients. Kaplan-Meier estimator survival analysis was performed in GlioVis.

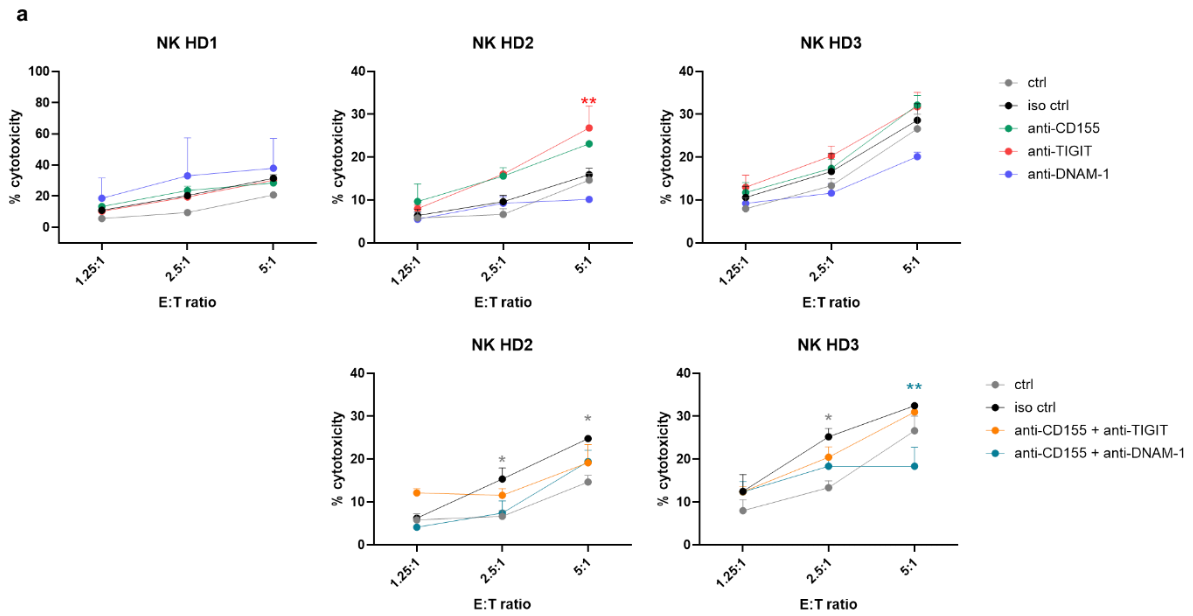


Figure S16: **The role of the CD155 axis in regulation of cytotoxicity of HD NK cells against NIB140 cells.** Shown are the results of calcein release assays performed with NK cells from three different healthy donors in which CD155, TIGIT, and DNAM-1 or their combinations were blocked by antibodies. Data are shown as mean ± SEM of three technical replicates. 2-way ANOVA with Tukey's multiple comparisons test was used for statistical analysis.

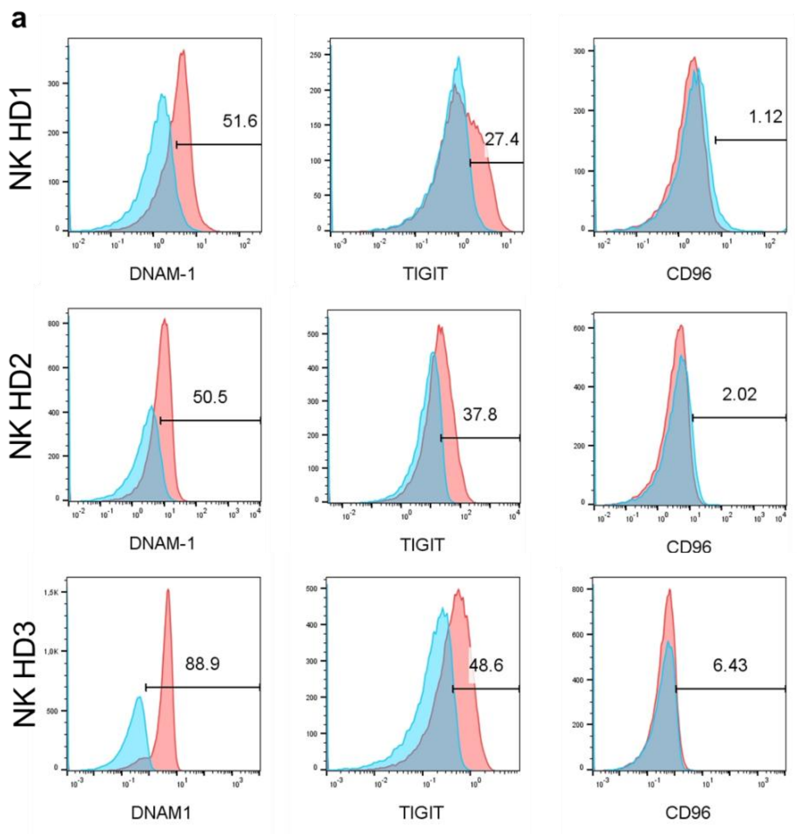


Figure S17: **Expression of DNAM-1, TIGIT and CD96 receptors on HD NK cells**. Shown are the results from the three donors that were used in calcein release assay to assess the effects of CD155, TIGIT and DNAM-1 blockade.
